# Density-dependent migration characteristics of cancer cells driven by pseudopod coordination

**DOI:** 10.1101/2021.11.04.467267

**Authors:** Gerhard A. Burger, Bob van de Water, Sylvia E. Le Dévédec, Joost B. Beltman

## Abstract

The ability of cancer cells to invade neighboring tissue from primary tumors is an important determinant of metastatic behavior. Quantification of cell migration characteristics such as migration speed and persistence helps to understand the requirements for such invasiveness. One factor that may influence invasion is how local tumor cell density shapes cell migration characteristics, which we here investigate with a combined experimental and computational modeling approach. First, we generated and analyzed time-lapse imaging data on two aggressive Triple-Negative Breast Cancer (TNBC) cell lines, HCC38 and Hs578T, during 2D migration assays at various cell densities. HCC38 cells exhibited a counter-intuitive increase in speed and persistence with increasing density, whereas Hs578T did not exhibit such an increase. Moreover, HCC38 cells exhibited strong cluster formation with active pseudopod-driven migration, especially at low densities, whereas Hs578T cells maintained a dispersed positioning. In order to obtain a mechanistic understanding of the density-dependent cell migration characteristics and cluster formation, we developed realistic spatial simulations using a Cellular Potts Model (CPM) with an explicit description of pseudopod dynamics. Model analysis demonstrated that strong coordination between pseudopods within single cells could explain the experimentally observed increase in speed and persistence with increasing density in HCC38 cells. Thus, the density-dependent migratory behavior could be an emergent property of single-cell characteristics without the need for additional mechanisms. This implies that coordination amongst pseudopods may play a role in the aggressive nature of cancers through mediating dispersal.

## Introduction

Breast Cancer (BC) is the most common cancer in women, and one of the main contributors to cancer mortality (WCRF, 2021). The primary cause of cancer mortality is metastasis, yet, because of its complexity, metastasis remains poorly understood (Fares et al., 2020; Suhail et al., 2019). Migration of cancer cells plays a crucial role in the metastatic cascade, not only for the long-range translocation of cells from the primary tumor to potential metastatic sites but also for the short-range dispersal of cells within the tumor, thus allowing accelerated tumor growth (Waclaw et al., 2015; Gallaher, Brown, and Anderson, 2019). A detailed understanding of cancer cell migration is essential to obtain insight into cancer progression and metastasis (Stuelten, Parent, and Montell, 2018), especially since expression of genes associated with migration is strongly associated with breast cancer survival (Nair et al., 2019).

A complicating factor in studying cancer cell migration is that BC is a highly heterogeneous disease. Two methods that are used to subdivide BCs into clinically-relevant subtypes are gene expression profiling and hormone receptor status (Viale, 2012). PAM50, a 50-gene classifier, divides BC into five intrinsic subtypes: luminal A, luminal B, basal-like, HER2 (human epidermal growth factor receptor 2)-enriched, and normal-like (Parker et al., 2009; Perou et al., 2000; Sørlie et al., 2001). This classification largely corresponds to classification by estrogen receptor (ER), progesterone receptor (PR), and HER2 status. BCs that are negative for these three receptors are called Triple-Negative Breast Cancers (TNBCs). Claudin-low BC, recently redefined as a BC phenotype rather than intrinsic subtype (Fougner et al., 2020), is characterized by its enrichment for Epithelial-Mesenchymal Transition (EMT) markers and stem cell-like features (Prat, Parker, et al., 2010).

Different (breast) cancer cells display a stunning variety in migratory strategies, and various methods have been developed to study these strategies *in vitro* (Kramer et al., 2013; Pijuan et al., 2019), *in vivo* (Beerling et al., 2016), and *in silico* (András Szabó and Merks, 2013; Wu et al., 2014; Huang et al., 2015; Te Boekhorst, Preziosi, and Friedl, 2016). Collective cell migration, where cells migrate in loosely or closely associated clusters, has been extensively studied in morphogenesis (Mayor and Etienne-Manneville, 2016), yet is also highly relevant during cancer metastasis (Rørth, 2009; Friedl et al., 2012). For example, in recent years the existence of intermediate EMT phenotypes is increasingly recognized. Such phenotypes are associated with the collective migration of tumor cell clusters (Brabletz et al., 2018), which can have 23-50 fold increased metastatic potential compared to single cells (Aceto et al., 2014). Despite this attention, many open questions remain regarding the mechanisms at play in such ‘social contexts’ (Angelini et al., 2011; Vedel et al., 2013). Recently, Jayatilaka et al. (2017) presented experimental evidence that paracrine IL-6/8 signaling amplified by cell density causes fast migration of MDA-MB-231 BC cells. Another approach was taken by Vedel et al. (2013), who studied the social context of 3T3 fibroblast cells using computational modeling, thereby demonstrating how complex collective migratory behavior can be an emergent property of single-cell migration properties. Thus, computational modeling is an invaluable tool to understand experimentally observed cell migration behavior, as hypothesized underlying mechanisms can be studied both at the single and collective level (Te Boekhorst, Preziosi, and Friedl, 2016). Various computational model formalisms for cell migration exist (as reviewed by Van Liedekerke et al., 2015; Te Bkohorst, Preziosi, and Friedl, 2016; Buttenschön and Edelstein-Keshet, 2020). The Cellular Potts Model (CPM) (Graner and Glazier, 1992; Glazier and Graner, 1993) is widely used for this purpose owing to its explicit incorporation of cell shape and flexibility to describe various biomechanical properties (András Szabó and Merks, 2013).

Here we study the migratory behavior of HCC38 and Hs578T, two highly migratory and invasive, claudin-low, basal B TNBC cell lines (Herschkowitz et al., 2007; Prat, Karginova, et al., 2013; Neve et al., 2006; Kao et al., 2009), using time-lapse microscopy and computational modeling with the CPM. To investigate how these cells respond to different social contexts, we plated these cells at various densities and performed 2D cell migration assays using Differential Interference Contrast (DIC) and fluorescence microscopy. In this setting, cell density clearly affected cell migration characteristics such as clustering, speed, and persistence for HCC38 cells, yet not for Hs578T cells. Specifically, at low densities HCC38 cells formed tight clusters, which loosened at high densities; this coincided with increased speed and persistence. We could not reproduce these density effects with published CPM models describing persistent cell migration, yet an extension of a CPM model of pseudopod-driven persistence with strong pseudopod coordination was sufficient to achieve the experimentally observed speed and persistence increase for HCC38 cells. Thus, pseudopodial activity can explain speed and persistence increase with density, provided that the pseudopods of a cell have the ability to affect each other’s extension.

## Results

### HCC38 and Hs578T cell lines both form streams during in vitro imaging

To investigate the migratory behavior of the TNBC cell lines HCC38 and Hs578T, we plated these lines in triplicate at 4 different cell densities within 24-well plates (20000, 50000, 100000, and 150000 cells per well). Subsequently, we performed a Random Cell Migration (RCM) assay (Roosmalen et al., 2011) using DIC and fluorescence microscopy of Hoechst-stained cells, imaged every 11-13 minutes for approximately 15 hours (Fig. S1, and Vid. S1). To quantify the migratory behavior of cells we performed automated cell tracking (Fig. 1A, Materials & Methods), by first segmenting the nuclei using Watershed Masked Clustering (WMC) (Yan and Verbeek, 2012) and then tracking the segmented nuclei in CellProfiler (Carpenter et al., 2006) using overlap tracking. Because of vignetting following stitching of adjacent images (see Fig. 1A DIC + Hoechst) and the high densities of cells in some fields of view (Fig. S1), some segmentation errors still occurred. Since these can affect the quantification of migration characteristics such as cell speed (J. B. Beltman, Marée, and Boer, 2009), we compared our automated tracking to manually determined tracks in a subset of wells. Analysis of the two methods of tracking revealed that they resulted in similar estimates for cell speed yet that automated tracking led to slightly lower instantaneous cell speeds compared to manual tracking (Fig. 1B). This minor difference could be explained by an overestimation of cell speed due to variability in manual center-of-mass determination (Huth et al., 2010). Therefore, and because overall differences between wells were similar for the two tracking approaches, we continued our analysis using automated tracking.

**Figure 1:**
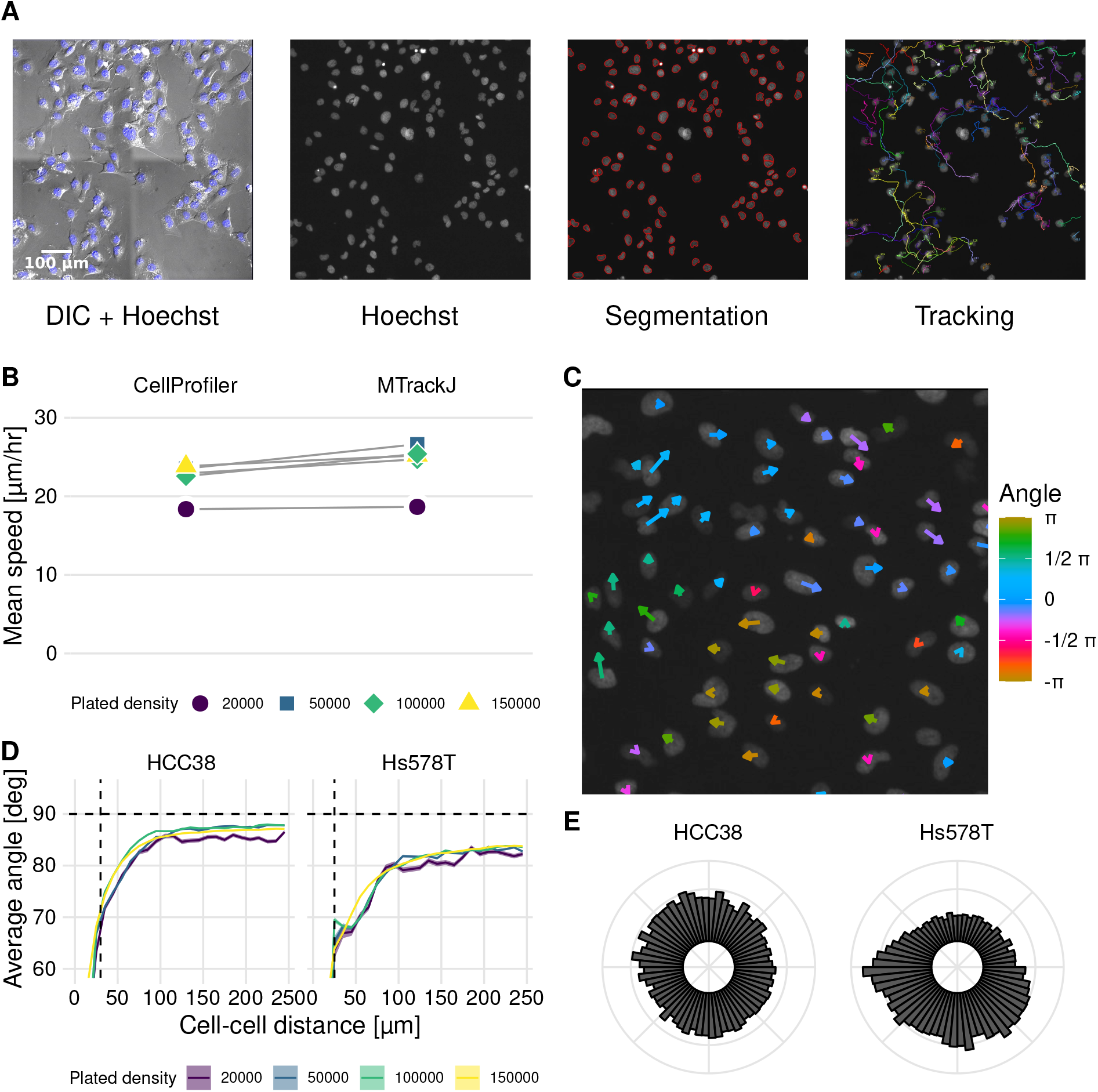
Identification of streams in automatically tracked videos of HCC38 and Hs578T. (A) Experimental setup and tracking workflow: nuclei were segmented using Hoechst, after which they were tracked. Images show HCC38 cells at density 50000 (frame 50 out of 71 frames). (B) Measured speed from automated tracking using CellProfiler and from manual tracking using MTrackJ. (C) Local migration directions at one time point in Hs578T cells. Size and color indicate instantaneous speed and current direction of migration. Image is a magnification of the top left corner of the bottom-rightmost video in S1 (Hs578T, 50000 cells, frame 51/71). (D) Analysis of migration angles between cell pairs as a function of the shortest distance between their centers of mass, at indicated plated cell densities. Horizontal dashed lines show theoretically expected average angle for random migration, vertical dashed lines show approximate nuclei diameter. (E) Polar histogram of migration directions of HCC38 and Hs578T cells.

A particularly striking feature that can be appreciated from the time-lapse videos is that Hs578T cells form ‘streams’ (Vid. S1, lower right; clockwise flow in Fig. 1C). To quantify this streaming behavior we analyzed the migration directions of all cells compared to the directions of all other cells. If cells were to migrate randomly, the average angle between two cell migration directions should approach 90 degrees (J. B. Beltman, Marée, and Boer, 2009). Consistent with the observation of streams within the videos, close-by cells had a lower average angle between their migration directions than remote cells (Fig. 1D). This streaming effect was more pronounced for Hs578T cells than for HCC38 cells and occurred at all densities, although visually it was mainly apparent at high densities. For both cell lines, but especially for Hs578T, the average angle remained below 90 degrees at all densities, which suggests a preferred direction of migration within wells. We confirmed this finding by polar histograms of cell migration direction (Fig. 1E), showing that the migration directions of HCC38 cells were approximately uniformly distributed, whereas Hs578T displayed a clear bias in migration direction. However, such large-scale streams could also be due to stage drift (J. B. Beltman, Marée, and Boer, 2009). Therefore, we took advantage of having two imaged locations per well (technical replicates), for which it would be expected that the polar plots would look highly similar if the streaming effects were due to stage drift. The individual polar plots of two associated well locations frequently exhibited a directional bias for Hs578T cells, yet this bias was typically different between the locations (Fig. S2A), strongly suggesting that streaming was not due to stage drift. Besides the presence of large-scale streams, both cell lines exhibited strong local streams, as evident from the strong decrease of migration angles for nearby cell movements compared to remote movements (Fig. 1D). Moreover, this difference in angle profiles remained present after correction for large-scale streams by subtracting the net overall displacement from each frame (Fig. S2B). In conclusion, both HCC38 and Hs578T cells formed local streams at all observed cell densities, and especially for Hs578T these streams occurred at a scale beyond the employed image dimensions.

### HCC38 cells form dynamic clusters at low densities

Visual inspection of images of HCC38 and Hs578T cells (Vid. S1) revealed that at low density HCC38 cells formed clusters (Fig. 2A top left), whereas Hs578T cells were less closely apposed to each other, although they may still be in contact via extended pseudopods (Fig. 2A top left). At high densities (Fig. 2A top right) clustering became less dominant for HCC38, yet remained similar for Hs578T. This clustering is surprising since HCC38 is a basal B cell line which are typically considered mesenchymal-like because of their high Vimentin levels (Fig. 2B). Consistent with differential clustering between the two cell lines at low density, Hs578T cells traveled further than HCC38 cells, as visible from tracks with starting points normalized to the origin (Fig. 2C). Nevertheless, the adhesion presumably driving HCC38 clustering did not completely prevent them from escaping these clusters, a feature that seems to be mediated by pseudopod-driven attachment (Vid. S1).

**Figure 2:**
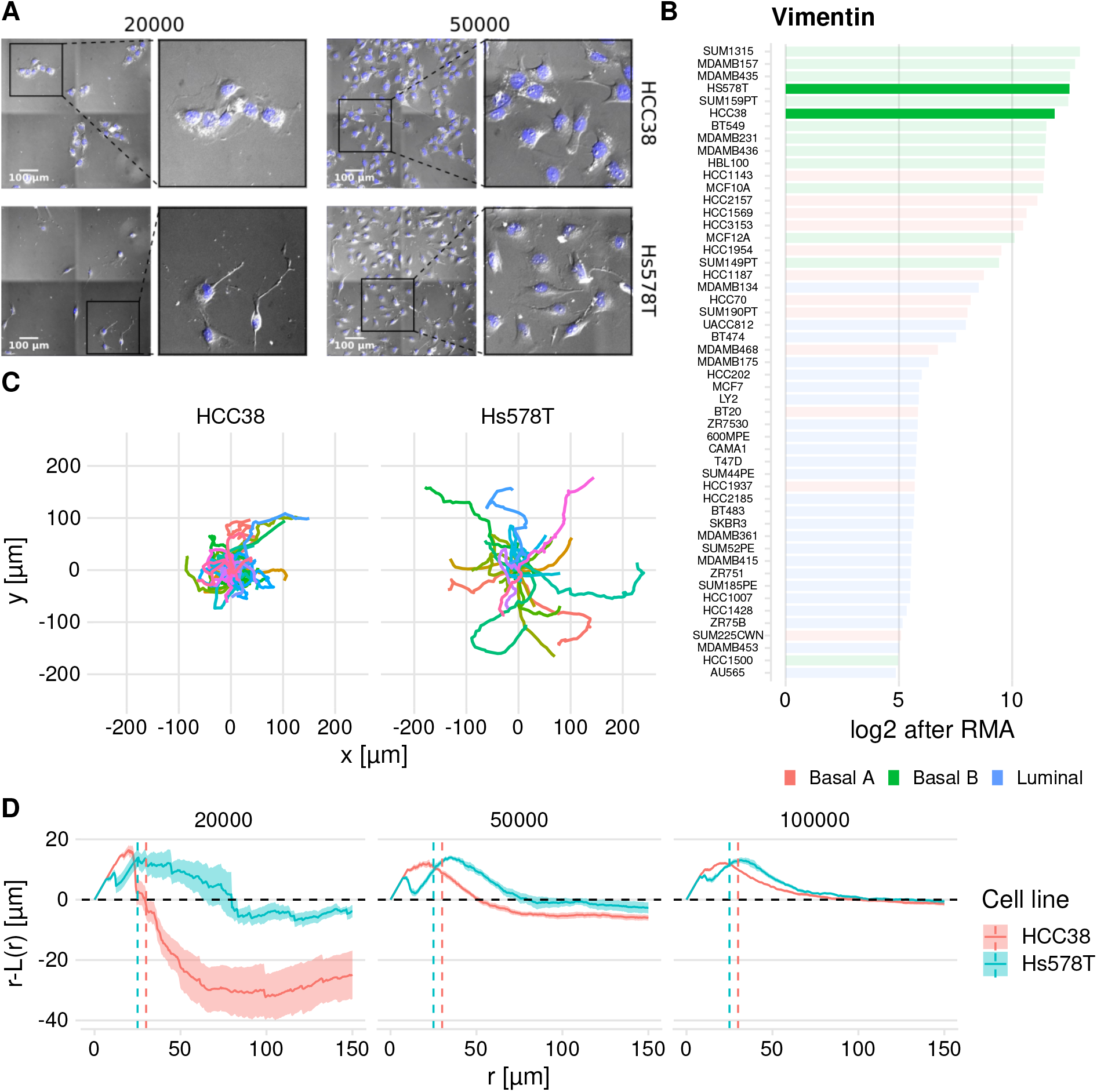
Quantification of observed dynamic clustering of HCC38 cells. (A) Still images of HCC38 (top row) and Hs578T (bottom row) at the two lowest densities at t ≈ 10 hours (insets show zoom-ins). (B) Vimentin expression of a collection of BC cell lines (data from Neve et al., 2006). Values are in log2 after Robust Multi-array Average (RMA). (C) Difference in maximum outreach of cells illustrated by normalized track plots (i.e. the starting point is moved to the origin). Data displayed from the same wells as Fig. 2A left column. (D) Spatial statistics visualization of HCC38 and Hs578T using the Ripley L function (see Materials & Methods) at t ≈ 10 hours. The dashed line *r* − *L*(*r*) = 0 shows the theoretically expected outcome in case of complete spatial randomness, values above and below this line signify dispersion and clustering. The vertical dashed lines denote approximate nuclei diameters.

To quantify clustering we employed spatial descriptive statistics by means of the Ripley L function (Ripley, 1977). Ripley L allows objective quantification of clustering compared to fully random placement of objects within a region, which is especially useful at high densities where differences in clustering are hard to determine by eye. Specifically, negative values for the quantity *r* − *L*(*r*) (see Materials & Methods) imply clustering, whereas positive values imply preferred dispersion. Beyond the diameter of an average nucleus (approximately 25 µm for Hs578T and 30 µm for HCC38), HCC38 cells clearly exhibited clustering, yet this clustering gradually disappeared at increasing densities (Fig. 2D, red). Hs578T cells did not cluster, but rather exhibited preferred dispersion (Fig. 2D, cyan), suggesting that these cells actively stay away from close apposition. Note that the initial increase in Fig. 2D shows short-range dispersion for both cell lines which is caused by volume exclusion (the small dips in this initial bump could be the result of occasional oversegmentation). Over time HCC38 clusters became less compact, as evident from an upward shift in the Ripley-L curves (Fig. S3). In conclusion, HCC38 cells formed clusters at low densities, whereas this was not the case for Hs578T cells.

### HCC38 cells exhibit increasing instantaneous speed and persistence with increasing density

Following our analysis of coordinated migration and clustering, we turned our attention to other aspects of cell migration, and whether these depended on cell density and cell type. First, we studied instantaneous speed, for which we investigated the relation with the observed cell densities within wells rather than with the plated densities (Fig. 3A), because these might differ due to spatial heterogeneity at different well locations. This speed analysis revealed that HCC38 cells move faster with increasing density (Fig. 3A left). In contrast, the speed of Hs578T cells was largely unaffected by cell density (Fig. 3A right), i.e., despite substantial experiment-to-experiment variability, there was a similar speed at all densities for each separate experiment.

**Figure 3:**
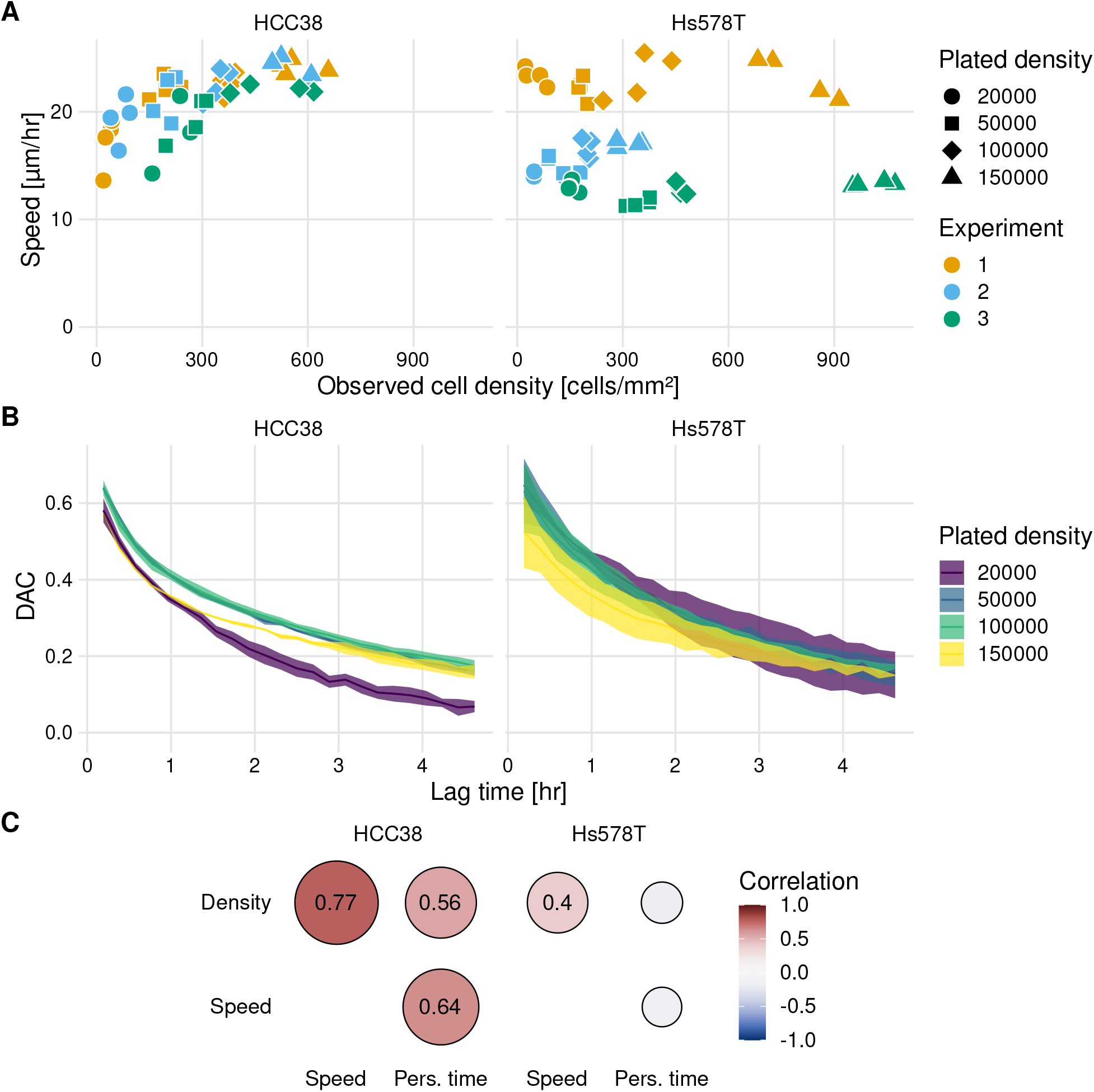
Analysis of the effect of cell density on cell speed and persistence. (A) Instantaneous speed for all individual wells as a function of the observed cell density within wells. Colors indicate results for the three separate experiments. (B) Mean ± SEM of the DAC as a function of the elapsed time for individual tracks. Note that the data for Hs578T from experiment 2 was excluded from this analysis because the differences in density were very small (see (A) and Fig. S4). (C) Correlogram showing the correlations between the observed cell density, instantaneous speed, and persistence time.

In addition to cell speed, a short-term measure of migration, we also analyzed the directional persistence of cells, a long-term measure of migration. A commonly used measure of persistence is the confinement ratio (also known as ‘straightness’ (Wortel, Dannenberg, et al., 2019), or ‘meandering index’ (Svensson et al., 2018)). However, since this measure is strongly biased by track duration (J. B. Beltman, Marée, and Boer, 2009; Gorelik and Gautreau, 2014) it is not suitable for our data which has substantial variation in track duration (Fig. S5). Instead, we analyzed persistence with Directional Auto Correlation (DAC) (Gorelik and Gautreau, 2014), which represents an unbiased measure of how fast cells lose their direction of migration (see Fig. 3a in Gorelik and Gautreau, 2014). The relation between the DAC and the lag time *τ*_lag_ can be characterized by the following exponential decay function:

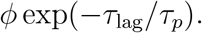

Here, *τ*_*p*_ is the persistence time and *ϕ* is the persistent fraction, a measure for the fraction of cells that is persistent (Vedel et al., 2013). Calibration of *τ*_*p*_ and *ϕ* for the two cell lines (Fig. S6) allowed us to study the correlations between cell density, speed, and persistence time. However, it should be noted that the optimal parameter fit did not always describe the decrease in the DAC well (see Fig. S7), so the resulting parameters should be interpreted cautiously, especially *ϕ*. For Hs578T, an increase in cell density was not associated with an effect on persistence time, matching earlier published results on 3T3 fibroblasts by Vedel et al. (2013). Interestingly, HCC38 cells reacted quite differently to increasing density: besides the strong positive correlation between cell density and speed, both cell density and speed also correlated with persistence time (Fig. 3C). In conclusion, HCC38 cells strongly increased their speed and persistence for increasing cell densities, whereas the Hs578T cell migration characteristics barely changed for increasing cell densities.

### Previous Cellular Potts migration models do not explain observed HCC38 speed-density behavior

Our observation that Hs578T cells exhibit dispersion rather than clustering or random positioning in space seems consistent with our findings that speed and persistence do not depend on cell density for this cell line: the cells essentially migrate as solitary cells at all densities. However, for HCC38 cells the relation between the observed clustering and the dependence of cell migration properties on density is less obvious. Therefore, we employed computational modeling to find the minimal requirements to achieve the observed density effects. A natural framework to model cell migration is the Cellular Potts Model (CPM), which is a grid-based formalism where multiple grid elements constitute a cell, and membrane elements stochastically extend and retract on the basis of a Hamiltonian. In the base CPM (see Materials & Methods) cell motion is driven only by adhesion and cell volume requirements, and therefore cells display Brownian motion, i.e. without any preferred direction and/or persistence (Wortel, Niculescu, et al., 2021). To model persistent cell migration realistically several extensions to the CPM have been proposed: 1) the ‘basic persistence model’ in which membrane extensions that move a cell in the same direction as its target direction (derived from previous moving directions) are favored (J. B. Beltman, Marée, Lynch, et al., 2007; Szabó et al., 2010), and 2) the Act model, which provides cells with persistence by modeling local actin dynamics (Niculescu, Textor, and Boer, 2015; Wortel, Niculescu, et al., 2021).

We explored a wide range of parameter values for both the basic persistence model and the Act model to investigate whether either of these CPM persistence extensions could reproduce the increases in speed and persistence with density observed in the HCC38 cell line. The basic persistence model does exhibit an increase in cell speed for increasing densities, yet there is no corresponding increase in persistence time (Fig. 4A). This finding goes against the universal coupling between speed and persistence that has been observed across many cell types in various *in vitro* and *in vivo* settings (Maiuri et al., 2015). Interestingly, the speed increase with density turns into a speed decrease with density when a connectivity constraint (Merks et al., 2006) is added that requires all membrane elements of a cell to remain in touch at all times, i.e., when cell mixing is hindered (Fig. S8 and Vid. S2). This suggests that the increase in speed depends on cells being able to mix. The Act-CPM model matches the observed data worse than the basic persistence model, exhibiting a decrease in both speed and persistence with increasing density (Fig. 4B and Vid. S3). Analysis of stream formation revealed that both CPM persistence models form local streams where cells align over a maximum of approximately three cell diameters (Fig. 4C). In conclusion, although published CPM extensions to describe cellular persistence lead to local streaming, these models are not consistent with the density dependence of HCC38 cell migration.

**Figure 4:**
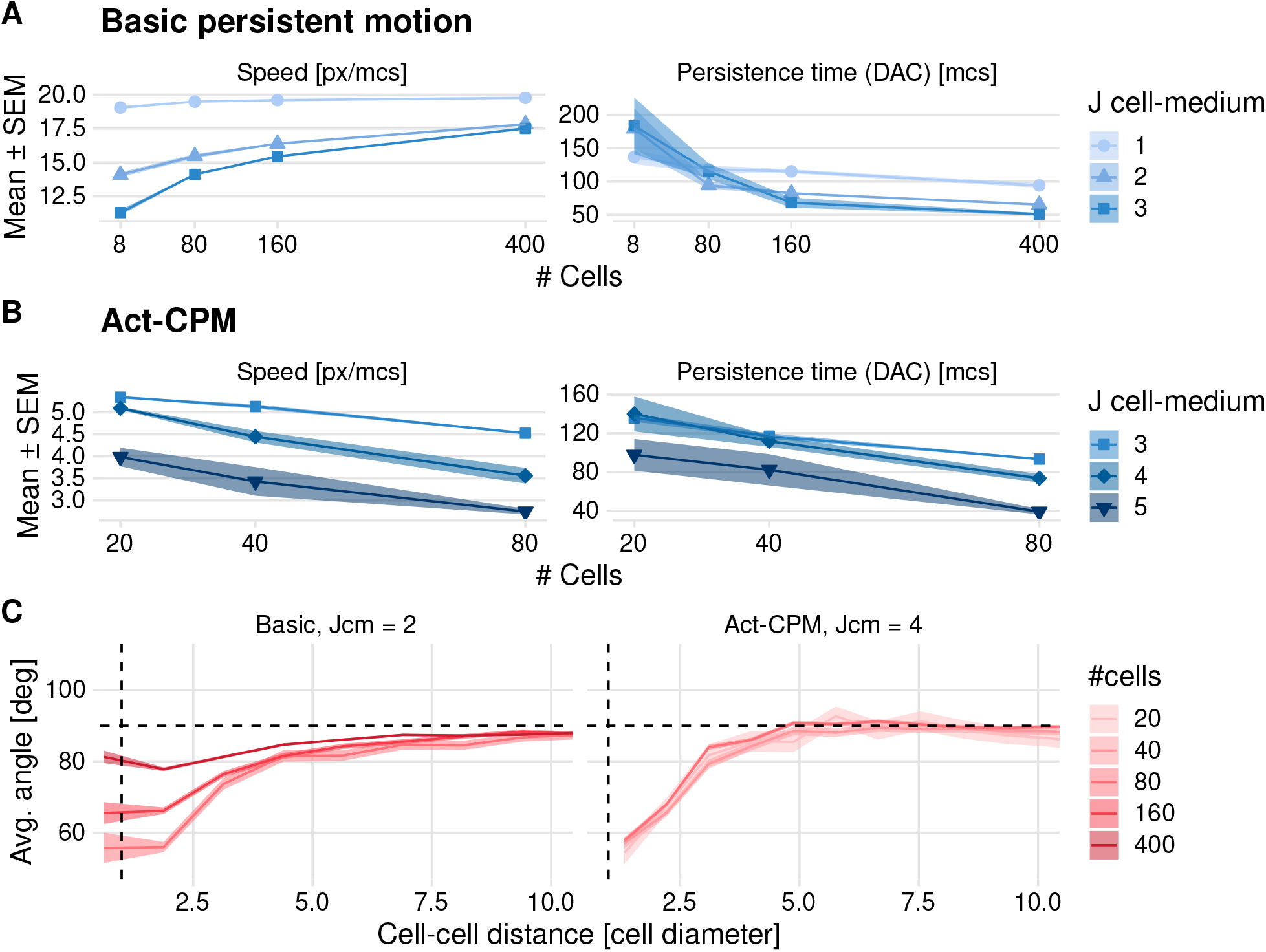
Effect of cell density on cell speed and persistence simulated using existing CPM persistence models. (A)-(B) Instantaneous speed and persistence time as a function of cell density for simulations with the basic persistence mechanism implemented in the Morpheus PersistentMotion plugin (A) and the Morpheus Act model implementation (B). Persistence time was in both cases fitted from the DAC in model simulations. (C) Stream formation in both models. Horizontal dashed lines denote theoretically expected average angle, vertical dashed lines denote approximate cell diameter.

### Coordinated pseudopod dynamics can explain density-dependent migratory behavior of HCC38 cells

Pseudopod formation is essential for persistent cell migration (Bergert et al., 2012; Van Haastert, 2011), and we observed high pseudopodial activity in the experimental videos, which seemed instrumental in HCC38 cells being able to move between clusters (Fig. 5A and Vid. S1). Cells simulated with the CPM extensions implementing either basic persistence or the Act model do exhibit a non-roundish shape with small extensions, but these are far shorter than the experimentally observed pseudopods, raising the question of whether the pseudopods might play a role in the observed density effects. Therefore, we implemented our previously developed CPM extension in Morpheus that was used to simulate the migration of dendritic-shaped tissue-resident memory T cells (Ariotti et al., 2012). In this model, cells form dendrite-like protrusions in the form of organized actin bundles that extend and retract within the cell and that move the cell in the direction of the protrusions (Fig. 5B). Although this model could achieve the dynamic clustering observed in HCC38 (Vid. S4), it could still not reproduce the experimentally observed dependence of speed and persistence on density (Fig. 5C). Similar to the Act model, the average speed decreased for increasing cell densities, presumably because the fixed dendrites obstruct each other’s extensions, thereby hampering (collective) migration, resulting in limited stream formation (Fig. 5D).

**Figure 5:**
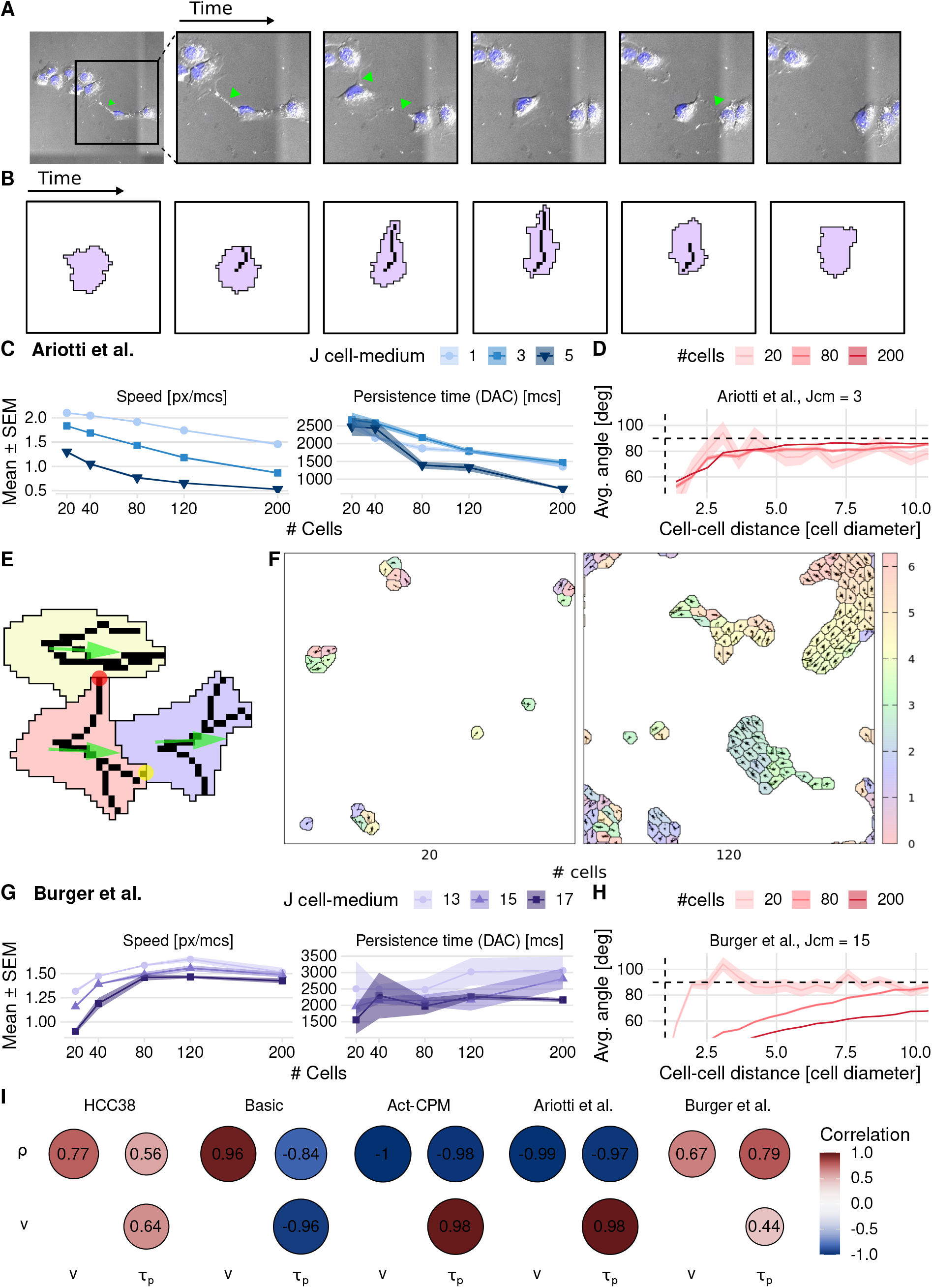
Effect of cell density on cell speed and persistence simulated using pseudopod-driven CPM persistence models. (A) Example of HCC38 cells at low density crossing a cluster using pseudopods (green arrows). (B) Illustration of how modeling protrusion/retraction of an actin fiber in a cell drives pseudopod-driven motility. (C) Results from the base pseudopod-driven model by Ariotti et al. (D) Stream formation in model by Ariotti et al. (E) Mechanisms added in the proposed model: adhesive pseudopod tips (yellow circle), Contact Inhibition of Locomotion-like pseudopod interaction (red circle), and pulling force in combined direction of the pseudopods (green arrows). (F) Snapshot of simulations using the proposed pseudopod-driven persistence model. Densities are comparable to HCC38 plated density of 20000 and 50000 (cf. Fig. S1). (G) Speed (left panel) and persistence (right panel) resulting from the proposed pseudopod-driven model. (H) Stream formation in the proposed pseudopod-driven model for different simulated cell numbers. (I) Correlograms comparing the different persistent models to the experimental correlations (*ρ*: density, *v*: speed, *τ*_*p*_: persistence time). Horizontal dashed lines in (D) and (H) denote theoretically expected average angle, vertical dashed lines show approximate cell diameter.

In order to test whether increased coordination between cellular pseudopods of a cell would matter for the dependence of migration on cell density, we adapted the modeled behavior of pseudopods in three ways (Fig. 5E; see Materials & Methods for details): First, we added an adhesive bonus to the pseudopod tips, because close observation of the experimental videos suggested that the pseudopods allow cells to attach to and pull on each other. Second, to further stimulate coordinated migration between cells, we implemented a type of Contact Inhibition of Locomotion (CIL) (Stramer and Mayor, 2017), where protrusions that are not aligned with the current overall movement direction of a cell and are touching other cells are quickly retracted and repolarized. Third, we let the pseudopods exert a pulling force to the cell as a whole in their combined direction (similar to Vedel et al., 2013). These three mechanisms together result in collective migration of clusters (Fig. 5F and Vid. S5). Importantly, our extended pseudopod model could explain the experimentally observed speed and persistence time increase with density for HCC38 cells (Fig. 5G, cf. Fig. S6). The simulated collective migration also goes hand-in-hand with cell alignment over whole clusters, which can be appreciated from the streaming quantification (Fig. 5H). Note though that in the simulations there is a direct dependence of the strength of the streaming on cell density, whereas in the experimental data this is not the case (cf. Fig. 1D). This cell-density dependence is less pronounced at lower surface energies *J*_cell,med_, for which there is also less long-range alignment (Fig. S9). In conclusion, our extended pseudopod model can explain an increase of speed and persistence time with increasing cell density as we observed for HCC38 cells, where other CPM migration models cannot (Fig. 5I). Strong coordination of pseudopod behavior within cells is thus an attractive explanation for the altered migration patterns with density.

## Discussion & Conclusion

TNBC is an aggressive subtype of breast cancer for which targeted therapies are just recently showing some promise (K. E. McCann, Hurvitz, and McAndrew, 2019). Since migration plays a crucial role in the metastatic cascade, more insight into the mechanisms behind TNBC migratory behavior could help identify potential targets for therapeutic intervention. Here we have used a combination of time-lapse microscopy and computational modeling to unravel the migratory behavior of HCC38 and Hs578T, two highly migratory TNBC cell lines. Both cell lines formed streams in our *in vitro* set-up, yet this was most clear from visual inspection in Hs578T cells. HCC38 cells formed dynamic clusters at low density which became less cohesive at high densities. Furthermore, HCC38 cells exhibited an increase in both speed and persistence time with increasing density. We could not reproduce this density dependence with CPM simulations implementing previously published persistence models, but a pseudopod-driven persistence model with strong coordination between the pseudopods of a cell could reproduce the key features of the experimentally observed HCC38 migratory behavior.

Given that HCC38 is a basal B TNBC cell line with very high Vimentin expression (Fig. 2B), one would expect that HCC38 is a mesenchymal cell-line (mentioned as such by Hollestelle et al., 2010; Kim et al., 2019). Thus, it was surprising that HCC38 cells strongly clustered, which is typically indicative of an intermediate EMT phenotype (Bocci, Kumar Jolly, and Onuchic, 2019). A possible explanation is that HCC38 is composed of epithelial and mesenchymal cells at a fixed ratio (as reported by Yamamoto et al., 2017). However, we could not identify subpopulations in our images, nor was this obvious in our single-cell migration analysis. Two indications that HCC38 is in fact a hybrid epithelial/mesenchymal cell line are that HCC38 has (1) high P-Cadherin expression (Kao et al., 2009), indicative of an intermediate EMT phenotype (Ribeiro and Paredes, 2014), and (2) high EpCAM (Epithelial Cell Adhesion Marker) expression (Klijn et al., 2015; Koedoot et al., 2021). Especially the increased EpCAM seems relevant because it has been reported to trigger “the formation of dynamic actin-rich protrusions” (Guillemot et al., 2001). Moreover, following EpCAM overexpression cell interactions are reduced to “sporadic contacts, mainly involving filopodia-like structures” (Litvinov et al., 1997), a description that matches our HCC38 observations (Vid. S1, cf. Fig. 2 in Winter et al. (2003)). This suggests EpCAM could play an important role in shaping pseudopodial interactions between HCC38 cells, and thereby in their migration characteristics.

Computational modeling of pseudopod-driven motility is a long-standing challenge (Schindler et al., 2021), and the incorporation of appropriate pseudopod mechanics in our CPM simulations was not straightforward. For example, based on the experimental images we aimed for long, finger-like extensions; however, for an approximately constant cell volume, such long pseudopods easily pull a cell apart in the CPM. One solution could be to use a compartmentalized CPM, with a separate nucleus and cytoplasm (Scianna and Preziosi, 2021). However, other model formalisms incorporating physical mechanisms in a spatially implicit (e.g., an Agent-Based Model (ABM) as applied in Vedel et al. (2013)), also represent an appropriate way to model coordinated behaviors of pseudopods. Our finding that HCC38 cells increase their speed and persistence with increasing cell density is somewhat exceptional. Earlier studies have usually reported cell speed to decrease (Angelini et al., 2011) or stay the same (C. P. McCann et al., 2010; Vedel et al., 2013) with increasing density. However, recently it has been shown that in MDA-MB-231, another claudin-low basal B TNBC cell line, paracrine IL-6/8 signaling amplified by cell density does cause faster migration for high than for low densities (Jayatilaka et al., 2017). Other examples of a cell-density-related speed increase include cell motion in endothelial monolayers (Szabó et al., 2010), or confined cell migration (Liu et al., 2015; András Szabó, Melchionda, et al., 2016). The contexts in which these other experiments were executed are somewhat different compared to our experimental setup, in which the speed increase occurred already at quite a low density (Fig. 3A). Nevertheless, it is possible that also in our experimental setting the observed density effects are (partially) due to a density-dependent nutrient gradient or chemotactic/chemokinetic signal. Still, we here showed that such density-dependent migration can occur as an emergent property of pseudopod coordination in single cells without any additional mechanisms.

Based on our simulations it seems that cells at low density can get ‘stuck’ in their respective clusters (Vid. S5), which is similar to the experimental observations (Vid. S1 top left). At high densities in our simulations, the clusters interact more, thereby avoiding rotating clusters, which causes an increase in persistence and speed. Nevertheless, at high densities the differences between simulations and experiments become more pronounced; whereas in the experiments the clusters became less cohesive, the simulations exhibit no difference with respect to cohesion (compare Fig. 1A HCC38 50000 with Fig. 5F 120). This is also reflected in the streams that form during simulations: Within the large migrating clusters that occur at high cell densities, cells’ migration directions become aligned over large distances (Fig. 5H). Lowering the surface energy between the cells and the medium *J*_cell,med_ results in less cohesive clusters, and a shorter-range alignment (Fig. S9). This suggests that cell adhesion might be decreased at high densities compared to low densities, which might also contribute to the high cell speeds at high density (see for example Fig. 5G).

In conclusion, in this study we shed light on the influence of cell density on the migratory behavior of two TNBC cell lines, HCC38 and Hs578T. We could reproduce the experimentally observed density-dependent speed increase in HCC38 cells using a pseudopod-driven CPM with strong coordination amongst pseudopods. A better understanding of the regulatory processes involved in pseudopod formation is urgently needed since they correlate with poor patient survival in multiple cancer types (Jacquemet et al., 2016). Our finding that strong pseudopod coordination can exacerbate the speed and persistence of cancer cells may be a partial explanation for the aggressive nature of such cancers due to high metastatic potential. Additionally, together with a previous report that showed how cell density affects the expression of cell-adhesion molecules (Stanley et al., 1995), the data presented here emphasize the need to include appropriate density-related controls in cell-migration assays.

## Materials & Methods

### Cell culture

Twenty-four hours prior to imaging, HCC38 (ATCC Cat# CRL-2314, RRID:CVCL_1267) and Hs578T (ATCC Cat# HTB-126, RRID:CVCL_0332) cells were seeded in complete medium on 24-well glass bottom plates (Sensoplate, Greiner Bio-One, 662892) coated with collagen (rat tail Type I, 10 µg mL^*−*1^), with the layout as shown in Table 1. The seeded densities were 20000, 50000, 100000, and 150000 cells per well, which, assuming uniform distribution in the well, corresponds to approximately 100, 250, 500, and 750 cells*/*mm^2^ One hour before imaging live Hoechst was added to the medium, and just before imaging the medium was refreshed (without additional Hoechst). The experiment was performed in triplicate.

**Table 1:** Plate layout for the random cell migration assays, the numbers denote the imaging order. There are two wells per condition, and two positions (technical replicates) per well. Wells were imaged by column in a zig-zag pattern.

|  | HCC38 |  | Hs578T |  | density |
| --- | --- | --- | --- | --- | --- |
|  | 1 | 2 | 3 | 4 |  |
| A | 1,2 | 15,16 | 17,18 | 31,32 | 20000 |
| B | 3,4 | 13,14 | 19,20 | 29,30 | 50000 |
| C | 5,6 | 11,12 | 21,22 | 27,28 | 100000 |
| D | 7,8 | 9,10 | 23,24 | 25,26 | 150000 |

### Microscopy

To allow nuclear tracking cells were incubated for one hour with Hoechst 33342. After incubation, the medium was refreshed and the plate was directly placed on an automated stage of a Nikon Eclipse TI equipped with a fluorescent lamp and 20x objective (Plan Apo, Air, numerical aperture (NA) 0.75, working distance (WD) 1.0), a Perfect Focus System (PFS) and a temperature- and CO_2_-controlled imaging chamber (custom design). Two positions per well were imaged using both fluorescence and DIC microscopy. The plates were imaged at 999 × 999 px (experiment 1) or 948 × 948 px (experiment 2 and 3) using a stitch of 2 × 2 positions with a pixel size of 0.79 µm. The imaging was repeated every 11 minutes (experiment 1) or 13 minutes (experiment 2 and 3) for 15 hours. Images are available at 10.5281/zenodo.5607734 (Le Dévédec, 2021)

Upon visual inspection of the microscopic images, we noted that for the third experiment of the HCC38 150000 condition cells were dying, therefore, we excluded these wells from further analysis.

### Image processing

The imaging processing and analysis consisted of multiple steps. Initially, proprietary Nikon ND2 image files were converted to the Tagged Image File Format (TIFF) using NIS-Elements (NIS-Elements, RRID:SCR_014329).

#### Automated tracking

Subsequently, the resulting TIFFs were processed in a CellProfiler pipeline (CellProfiler Image Analysis Software, V2.1.1, RRID:SCR_007358) (Carpenter et al., 2006), containing the following steps:

- Cropping: Following stitching of the images, they contained zero-intensity patches at the edges as a result of (mis)alignment. To avoid problems with segmentation and edge detection later in the pipeline, we cropped the images by 2 pixels at the edges.
- Segmentation: The cropped images were segmented using the WMC approach (Yan and Verbeek, 2012). See Table 2 for the utilized parameters.
- Object identification: We converted the connected components in the segmented images into objects. The resulting objects were filtered on size; we only retained objects with a diameter between 10 px (8 µm) and 40 px (32 µm) for Hs578T, or 50 px (40 µm) for HCC38. Additionally, we discarded objects touching the image border to prevent inaccurate center-of-mass calculation.
- Tracking: We tracked the remaining objects using the *Overlap* tracking method with a maximal pixel distance of 30.

#### Manual tracking

To compare our automated tracking to manual tracking, we used MTrackJ (Meijering, Dzyubachyk, and Smal, 2012) in Fiji (Fiji, RRID:SCR_002285) (Schindelin et al., 2012; Rueden et al., 2017) to manually track a representative subset of the wells by clicking the center of mass of each cell in each frame. Although manual tracking is considered the gold standard for tracking (Cordelières et al., 2013), variability in center-of-mass determination (e.g. due to operator fatigue) can cause an overestimation of the actual cell speed (Huth et al., 2010).

**Table 2:** Settings for the WMC plugin in CellProfiler (see Yan and Verbeek, 2012, for further details).

|  | HCC38 | Hs578T |
| --- | --- | --- |
| Gaussian filter size | 2 | 1 |
| Rolling ball size | 200 |  |
| Use parabolic kernel | No |  |
| Noise | 15 | 12 |
| Low seed | 15 |  |
| High seed | 20 |  |
| Low bound | 0.4 |  |
| High bound | 0.7 |  |
| Use intensity equalize | No |  |

#### Nucleus diameter calculation

Because the cells show high pseudopodal activity the cell diameters are difficult to estimate. Instead, we estimated the nucleus diameters, which are 30 µm and 25 µm for HCC38 and Hs578T. Using the EBImage R package (Pau et al., 2010), we first applied an adaptive threshold on the Hoechst signal, followed by watershed transformation and object feature analysis. Nucleus diameters were estimated as two times the average nuclei radius reported by EBImage, rounded up. These nucleus diameters serve as an approximation for the nearest possible distance between cells.

### Track Analysis

Tracking data from CellProfiler was imported into R using an in-house developed script (Wink and Burger, 2021, Wink, 2015, Ch. 7), and by fixing t rack identifiers wi th the CPTrackR package (Burger, Heldring, and Wink, 2021). MTrackJ data were imported using the mdftracks package (Burger, 2021).

Analysis in R (R Project for Statistical Computing, RRID:SCR_001905) (R Core Team, 2018) was performed with RStudio (RStudio, RRID:SCR_000432) (RStudio Team, 2016) and with the packages celltrackR (Wortel, Dannenberg, et al., 2019), spatstat (Baddeley, Rubak, and Turner, 2015), and tidyverse (Wickham, 2017) packages.

### Directional Autocorrelation

The Directional Auto Correlation (DAC) of all cells was computed by

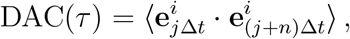

where 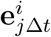 denotes the normalized direction of motion of cell *i* at time *j*Δ*t*, and the angle brackets denote averaging over all cells *i* and all times *j*Δ*t* and (*j* + *n*)Δ*t*, where Δ*t* is the sampling time, of which the lag time *τ* = *n*Δ*t* is a multiple. We computed the DAC in R using the overallNormDot function, which we contributed to the celltrackR package (Wortel, Dannenberg, et al., 2019).

After removing DAC(0), which is by definition equal to unity, we fitted the exponential decay function

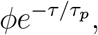

which gives an estimate for the weight factor *ϕ* (can be interpreted as the fraction of cells that is persistent) and persistence time *τ*_*p*_ (Vedel et al., 2013). Since the estimates for *ϕ* resulting from parameter calibration were not always reliable (see Fig. S7) we focussed on *τ*_*p*_ in our further analysis.

### Correlograms

Averaged correlation values for the correlograms were computed using the Fisher transformation. First, the Pearson correlation *r* for each experiment and simulation replicate was converted into a Fisher’s *z*:

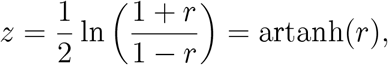

where artanh is the inverse hyperbolic tangent. Then *z* can be averaged and converted back with

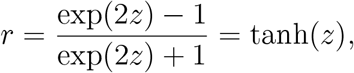

where tanh is the hyperbolic tangent (Corey, Dunlap, and Burke, 1998).

### Clustering analysis with Ripley K

To analyze spatial clustering we used the common transformation on Ripley K (Ripley, 1977) defined as

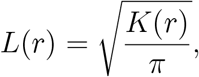

and provided in the spatstat R package (Baddeley, Rubak, and Turner, 2015). We subsequently visualized *r* − *L*(*r*) as a function of *r* such that in case of complete spatial randomness

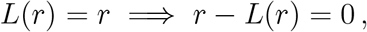

which allows to determine whether clustering (*r* − *L*(*r*) *<* 0) or dispersion (*r* − *L*(*r*) *>* 0) occurs.

### Cellular Potts Modeling

In the Cellular Potts Model (CPM) cells are defined as a collection of lattice sites **v** ∈ ℤ^*n*^ with the same cell identifier *σ*. Each cell also has an associated cell type *τ* (*σ*). The energy within cells (i.e., lattice sites having shared *σ*) is zero, but at sites forming the cell boundaries (referred to as ‘membrane elements’ below), there is a cell-type-dependent surface energy *J*_*τ1,τ*2_. A simulation consists of a sequence of Monte Carlo Steps (MCSs), during which cells attempt to extend membrane elements that would modify the cell identifier of a lattice site *σ*(**v**) into the identifier of one of the neighboring lattice sites *σ*(**v**_**n**_) in their 2D Moore neighborhood (i.e., 8 sites). The probability that such an extension is accepted depends on the change in the Hamiltonian:

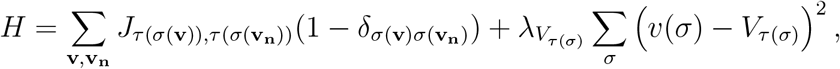

where the first term is the sum of the surface energies over all **v, v**_**n**_ neighbor pairs, and the second term is the elastic volume constraint which keeps cells within a range of biologically appropriate sizes. Furthermore, *δ* is the Kronecker delta, *λ*_*V*_τ is the elastic constant for the volume of cell type 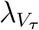 is the target volume of cell type *τ*, and *v*(*σ*) is the actual volume of cell *σ*.

The probability *p* that an extension is accepted depends on the change in the Hamiltonian Δ*H* as follows:

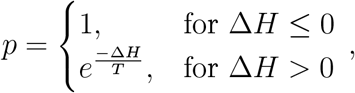

where *T* is the temperature (Graner and Glazier, 1992; Glazier and Graner, 1993). We used Morpheus (RRID:SCR_014975, Starruß et al., 2014) for the CPM implementation.

#### Existing persistence models

##### Basic persistence

In the basic persistence model implemented in the PersistentMotion plugin in Morpheus, cells have a target direction **t** based on previous movements which is updated continuously according to

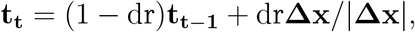

where dr = min(1*/*dt, 1) is the decay rate, with decay time dt in Monte Carlo Step (MCS), and **Δx** = **x**_**t**_ − **x**_**t***−***1**_ is the shift of the cell centroid in the previous MCS. For a proposed copy attempt *σ*(**v**) → *σ*(**v**_**n**_) in update direction **s**, the additional change in Hamiltonian *H* due to persistence is computed as:

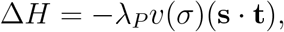

where *λ*_*P*_ is the strength of persistence, and *v*(*σ*) is the cell volume. For details on the parameters used for the basic persistence model see Table S1.

##### Act-CPM

In the Act-CPM model, persistence is achieved by recording each lattice site’s “actin activity”, which depends on the MCSs elapsed since its most recent protrusive activity. Upon a successful copy attempt, the target lattice site is assigned the maximum activity value (Max_act_), which decreases every MCS until it reaches zero. By making a copy attempt from an active site into a less active site more favorable, a local positive-feedback mechanism is created which causes persistent motion. For a proposed copy attempt *σ*(**v**) → *σ*(**v**_**n**_) the additional change in Hamiltonian *H* is computed as

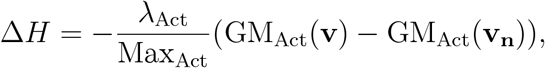

where GM_Act_(**v**) is the geometric mean of all activities of lattice sites in the Moore neighborhood of **v** which share the same cell identifier *σ*(**v**) (Niculescu, Textor, and Boer, 2015; Wortel, Niculescu, et al., 2021).

For this study we used the Act-CPM model provided in Morpheus, for details on the parameters used see Table S2.

#### Pseudopod dynamics

To obtain realistic pseudopod-driven persistence matching our experimental observations, we adapted a published model by (Ariotti et al., 2012). In this model, pseudopod dynamics are realized by extending explicitly described actin fibers from the center-of-mass of the cell in the direction obtained by drawing a value from a von Mises distribution centered around the current movement direction of the cell. This actin fiber can only grow inside the cell, and cell growth directly next to the actin fiber is promoted (via a *DeltaH* extension), and cell shrinking is prevented. After a maximum growth time the actin fiber is retracted (see SI Ariotti et al., 2012, for details)). See Fig. 5B for an example of this pseudopod-driven motility with a single pseudopod.

We implemented this model in a Morpheus plugin, and we adapted it in several ways to implement various processes involved in the coordination of pseudopod behavior:

- In HCC38 cells *in vitro* we observed a ‘stickiness’ of pseudopods (Vid. S1). To mimic this effect we added an adhesion bonus `tip-bonus` to the in silico pseudopod tips. The bonus is applied when a proposed cell extension is within the `max-distance-for-tip-bonus` of one of its pseudopods. Moreover, when this position is also within the `max-distance-for-tip-bonus` of a pseudopod from a neighboring cell, the bonus is doubled.
- To increase the persistence of the cells, we implemented a pseudopod pulling effect, similar to an effect simulated in Vedel et al. (2013)). Given a copy attempt *σ*(**v**) → *σ*(**v**_**n**_) in update direction **s**, the change in Hamiltonian is computed as

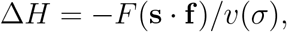

where *F* is the pulling force, **f** is the vector sum of all pseudopods, and *v*(*σ*) is the cell volume.
- To increase the alignment of neighboring cells, we implemented a ‘touch’ strategy inspired by Contact Inhibition of Locomotion (CIL). Specifically, when a pseudopod touches a neighboring cell it is immediately set to the retraction phase allowing pseudopod repolarization. This implementation is executed when the `touch-behavior` parameter is set to `retract`. Another touch strategy we implemented includes an additional rule when a pseudopod touches a neighboring cell laterally (i.e. when cos *α <* .85, with *α* the angle between the pseudopod direction and the current movement direction of the cell). In that case, the pseudopod is instantly retracted (i.e. skipping the retraction phase), leading to immediate repolarization of the pseudopod. Note that this implementation is executed when the `touch-behavior` parameter is set to `poof-dir`. The results presented here are obtained with the `poof-dir` touch strategy.

See Table S4 for a full list of parameters for the CPM pseudopod extension, and Table S5 for the values used. The code for the Morpheus plugin is available at 10.5281/zen-odo.5484491 (Burger and J. Beltman, 2021).

#### Choice of simulation parameters

To efficiently explore the parameter space, we used the Python Programming Language (RRID:SCR_008394) in Jupyter Notebook (RRID:SCR_018315) (Kluyver et al., 2016), and FitMultiCell (Alamoudi et al., n.d.) (based on pyABC (Schälte et al., 2021) and Morpheus (Starruß et al., 2014)). Based on this extensive exploration, we selected representative parameter sets that qualitatively matched (parts of) the experimental data. The simulated number of cells were similar as the number of cells observed in the experiments (Fig. S4), and all simulations ran for 20000 MCS.

### Simulation measurements

We saved the cell positions from the simulations every MCS, and after discarding of the first 1000 mcs to allow for equilibration, we analyzed them in the same way as the experimental cell tracks, except for instantaneous speed, which was estimated based on 50 mcs subtracks.

## Acknowledgments

We would like to thank the Center for Information Services and High Performance Computing, Technische Universität Dresden for providing the Morpheus modeling and simulation environment (Starruß et al., 2014; Morpheus, n.d.), and in particular Jörn Starruß and Walter de Back for their prompt and helpful communication. We also thank András Szabó for useful discussions. This work was supported by a Vidi grant from the Netherlands Organization for Scientific Research (NWO; grant 864.12.013 to JBB).

## Supplementary Material

### Materials & Methods

**Table S1:**
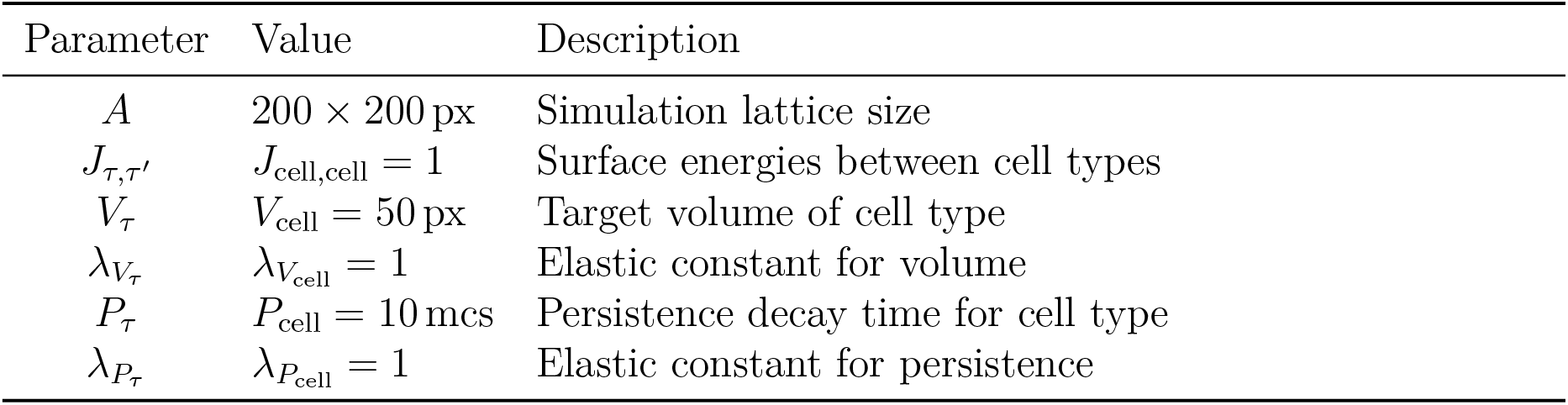
Parameters for the Morpheus PersistentMotion model used to generate Fig. 4A.

| Parameter | Value | Description |
| --- | --- | --- |
| $A$ | $200 \times 200$ px | Simulation lattice size |
| $J_{\tau,\tau'}$ | $J_{\text{cell,cell}} = 1$ | Surface energies between cell types |
| $V_{\tau}$ | $V_{\text{cell}} = 50$ px | Target volume of cell type |
| $\lambda_{V_{\tau}}$ | $\lambda_{V_{\text{cell}}} = 1$ | Elastic constant for volume |
| $P_{\tau}$ | $P_{\text{cell}} = 10$ mcs | Persistence decay time for cell type |
| $\lambda_{P_{\tau}}$ | $\lambda_{P_{\text{cell}}} = 1$ | Elastic constant for persistence |

**Table S2:**
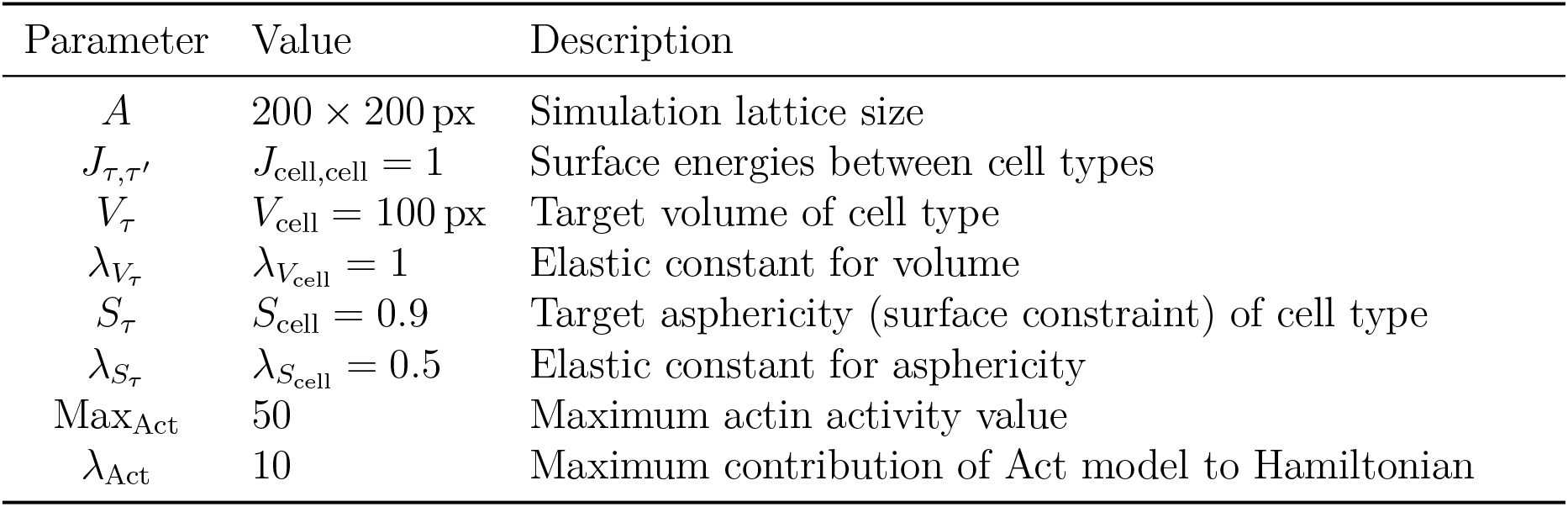
Parameters for the Morpheus Act-CPM plugin model used to generate Fig. 4B.

**Table S3:** Shared parameters for the models used to generate Figs. 5C and 5G.

| Parameter | Value | Description |
| --- | --- | --- |
| $A$ | $400 \times 400$ px | Simulation lattice size |
| $J_{\tau,\tau'}$ | $J_{\text{cell,cell}} = 1$ | Surface energies between cell types |
| $V_{\tau}$ | $V_{\text{cell}} = 250$ px | Target volume of cell type |
| $\lambda_{V_{\tau}}$ | $\lambda_{V_{\text{cell}}} = 1$ | Elastic constant for volume |
| $S_{\tau}$ | $S_{\text{cell}} = 0.9$ | Target asphericity of cell type |
| $\lambda_{S_{\tau}}$ | $\lambda_{S_{\text{cell}}} = 0.5$ | Elastic constant for asphericity |

**Table S4:** Parameter description for the developed Morpheus Pseudopodia plugin. **Bold parameters** are parameters introduced specifically for this study.

| Parameter | Description |
| --- | --- |
| field | Morpheus field to keep track of the actin cytoskeleton |
| moving-direction | The (buffered) moving direction of the cell (used to create new pseudopodia in that approximate direction) |
| max-growth-time | Max number of MCS pseudopods grow |
| max-pseudopods | Max number of pseudopods per cell |
| time-between-extensions | Refractory period for pseudopod extension |
| tip-bonus | **Adhesion bonus for the pseudopod tip applied to the Hamiltonian.** |
| max-distance-for-tip-bonus | **The maximum distance at which surrounding pixels are considered part of the pseudopod tip** |
| neighboring-actin-bonus | Hamiltonian bonus to stimulate growth directly next to actin cytoskeleton |
| init-dir-strength | $\kappa$ for von Mises distribution to bias pseudopod formation in moving-direction |
| cont-dir-strength | Same as init-dir-strength, except to bias pseudopod growth direction |
| retraction-mode | • backward: pseudopod is retracted backwards<br>• forward: pseudopod is “retracted” forwards<br>• in-moving-direction: forwards if aligned with moving-direction, backwards otherwise. |
| touch-behavior | **If pseudopods touch another cell...**<br>• nothing<br>• retract: retract with retraction-mode<br>• attach: stop pseudopod growth phase and begin delayed retraction phase<br>• poof-dir: if touching laterally, allow pseudopod to repolarize immediately |
| pull | **Turn ‘pulling’ on/off** |
| pull-strength | **Hamiltonian bonus if update moves cell in the combined direction of the pseudopods** |

**Table S5:** Parameters for Pseudopodia plugin used together with Table S3 to generate Figs. 5C and 5G. See Table S4 for a description of these parameters.

| Parameter | Ariotti | Burger |
| --- | --- | --- |
| max-growth-time | 20 |  |
| max-pseudopods | 3 |  |
| time-between-extensions | 1 |  |
| tip-bonus | 0 | 30 |
| max-distance-for-tip-bonus | n/a | 5 |
| neighboring-actin-bonus | 8 |  |
| init-dir-strength | 8 |  |
| cont-dir-strength | 16 |  |
| retraction-mode | forward |  |
| touch-behavior | nothing | poof-dir |
| pull | false | true |
| pull-strength | n/a | 50 |

### Figures

**Figure S1:**
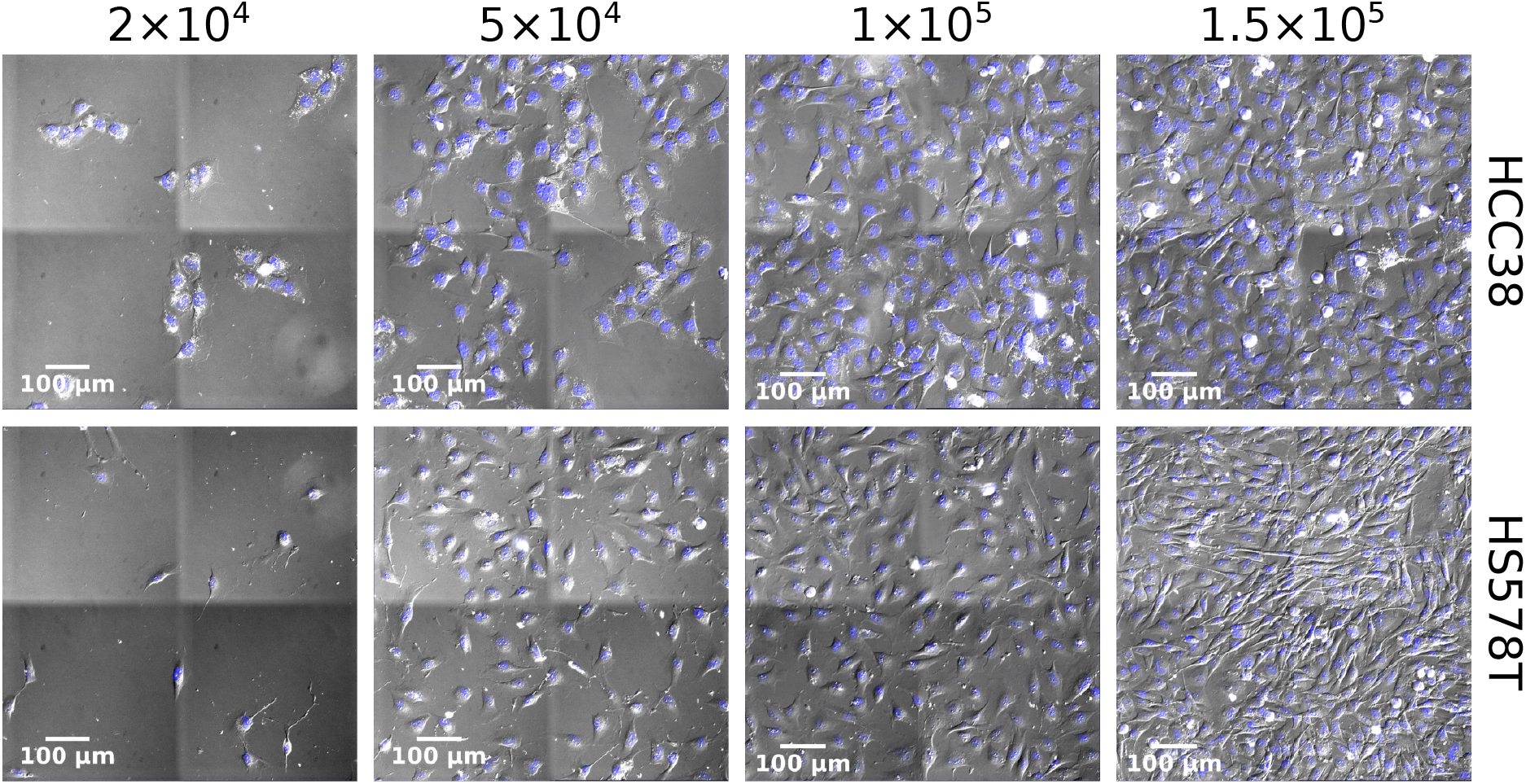
Overview of all experimental plated densities for HCC38 and Hs578T cell lines. See Vid. S1 for the corresponding videos.

**Figure S2:**
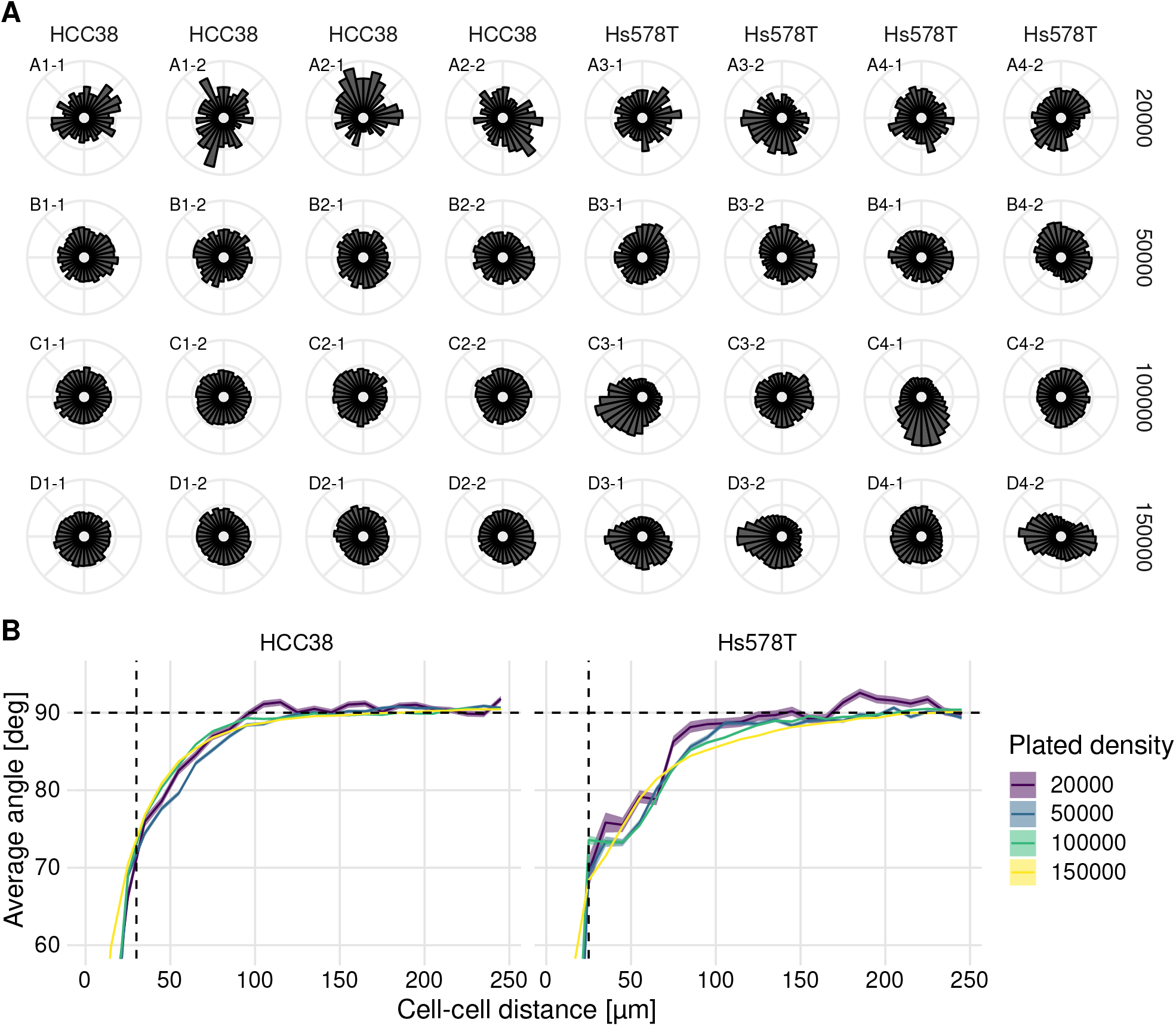
Analysis of large-scale and local streams. (A) Polar histogram of migration directions within a single experiment (24-well plate, see Table 1 for layout). Note that every 2 columns contain technical replicates from the same well, so any potential stage drift should have shown up in at least the technical replicates, but likely also in the whole plate. (B) Flow summary with drift correction (subtraction of net overall movement per frame). Horizontal dashed lines denote theoretically expected average angle, vertical dashed lines denote approximate cell diameter.

**Figure S3:**
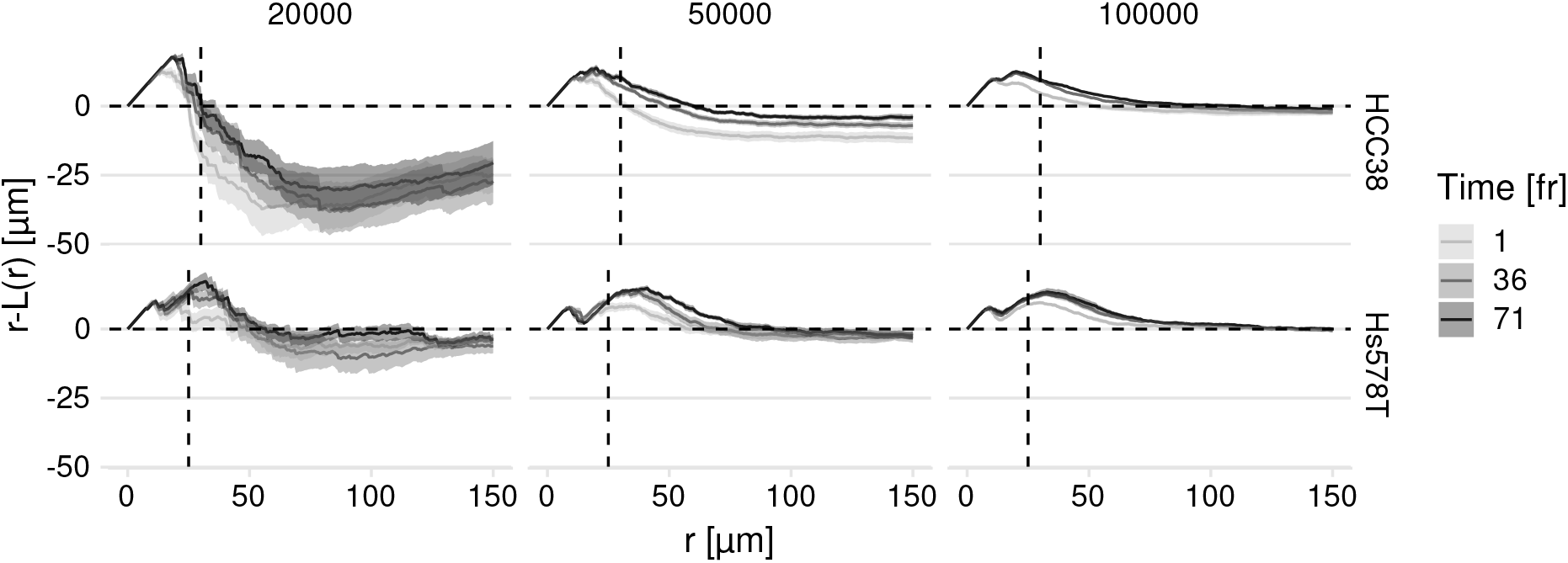
Decrease in clustering over time. The dashed line *r* − *L*(*r*) = 0 shows the theoretically expected outcome in case of complete spatial randomness, values above and below this line signify dispersion and clustering. The vertical dashed lines denote approximate cell diameters.

**Figure S4:**
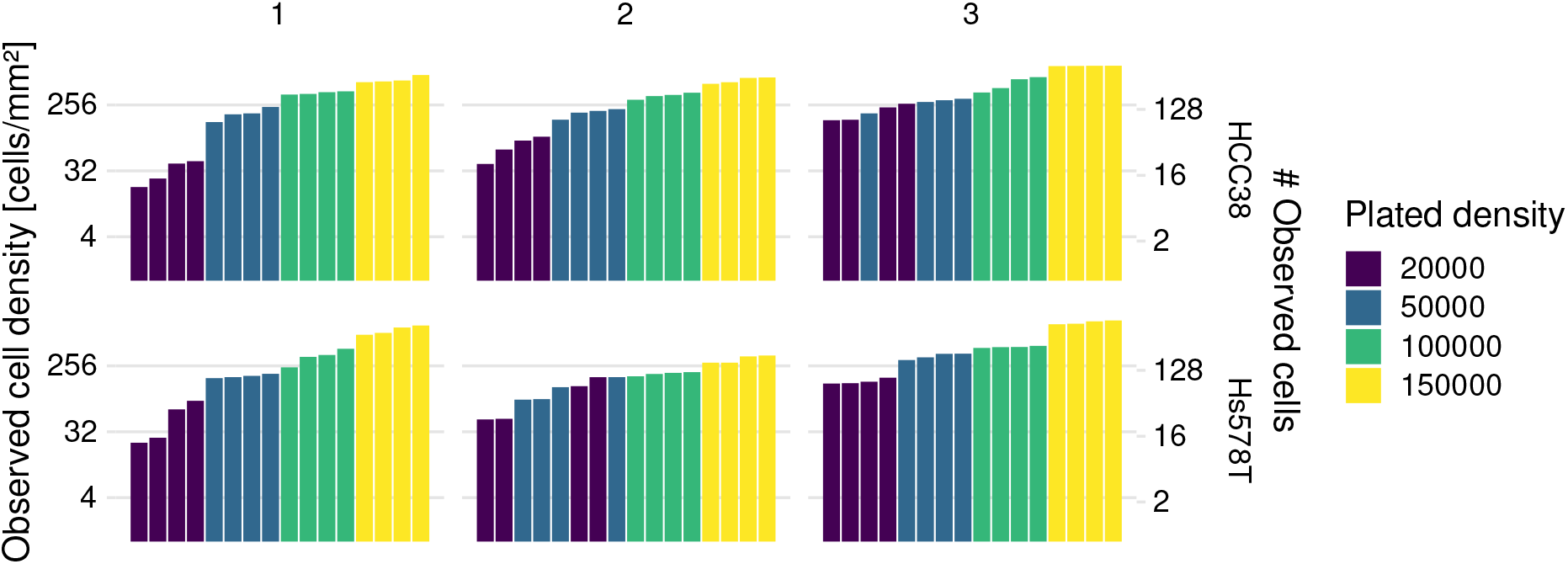
Plated density versus observed cell density in the images for each imaged position. There are 2 wells × 2 positions per condition, and positions are ranked on the number of observed cells. Note that the conditions in Hs578T experiment 2 are not separated well, which is why they were excluded from the DAC analysis summary in which plated density was used as independent variable (Fig. 3B).

**Figure S5:**
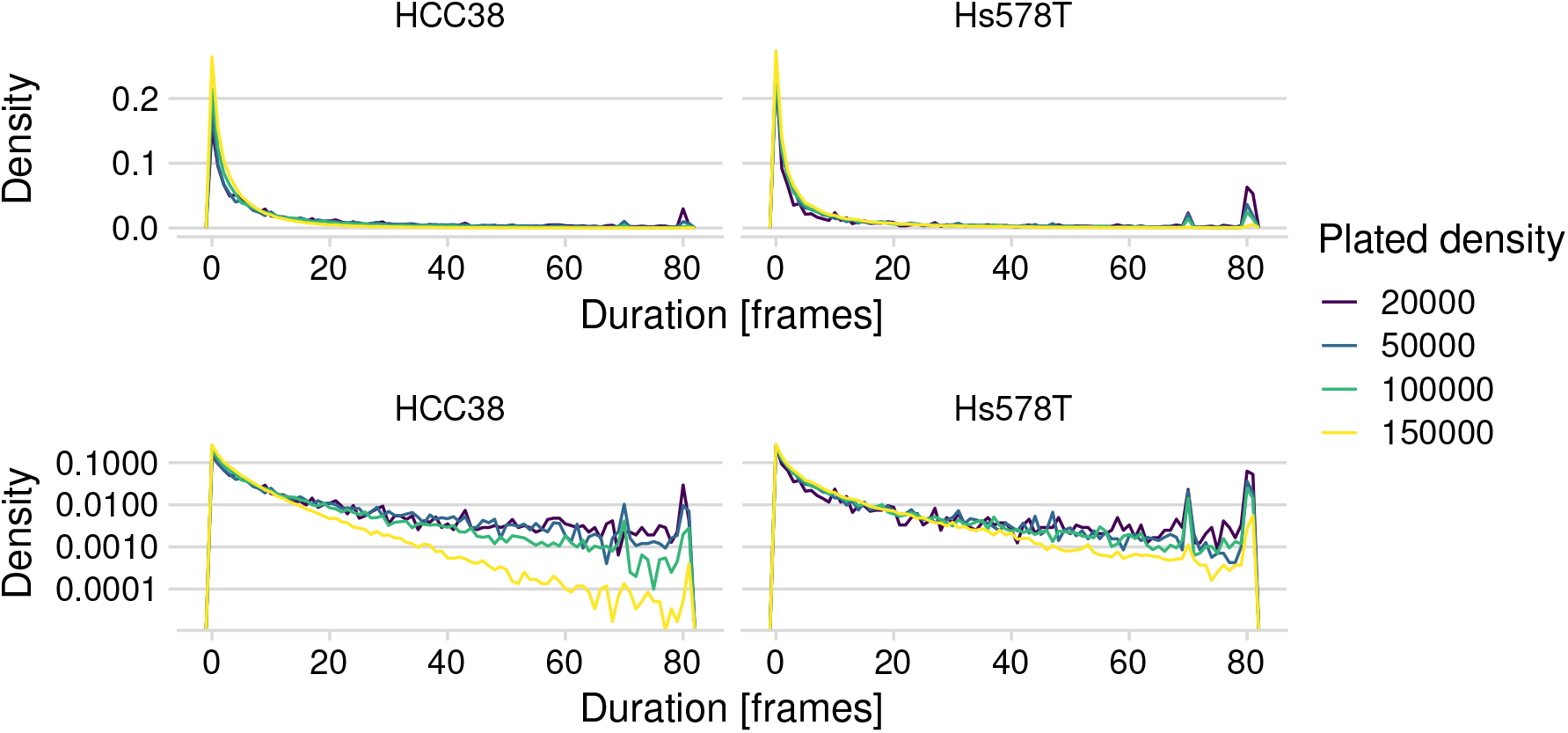
Density plot of track durations. The top panels have a linear vertical axis, and the bottom panels have a logarithmic one. Most tracks are only a few frames long, which is likely due to segmentation and conservative tracking. Especially for HCC38, there are relatively few cells that can be tracked for the complete duration of the experiment. Peaks at 71 and 81 mark the maximum track length for different experiments.

**Figure S6:**
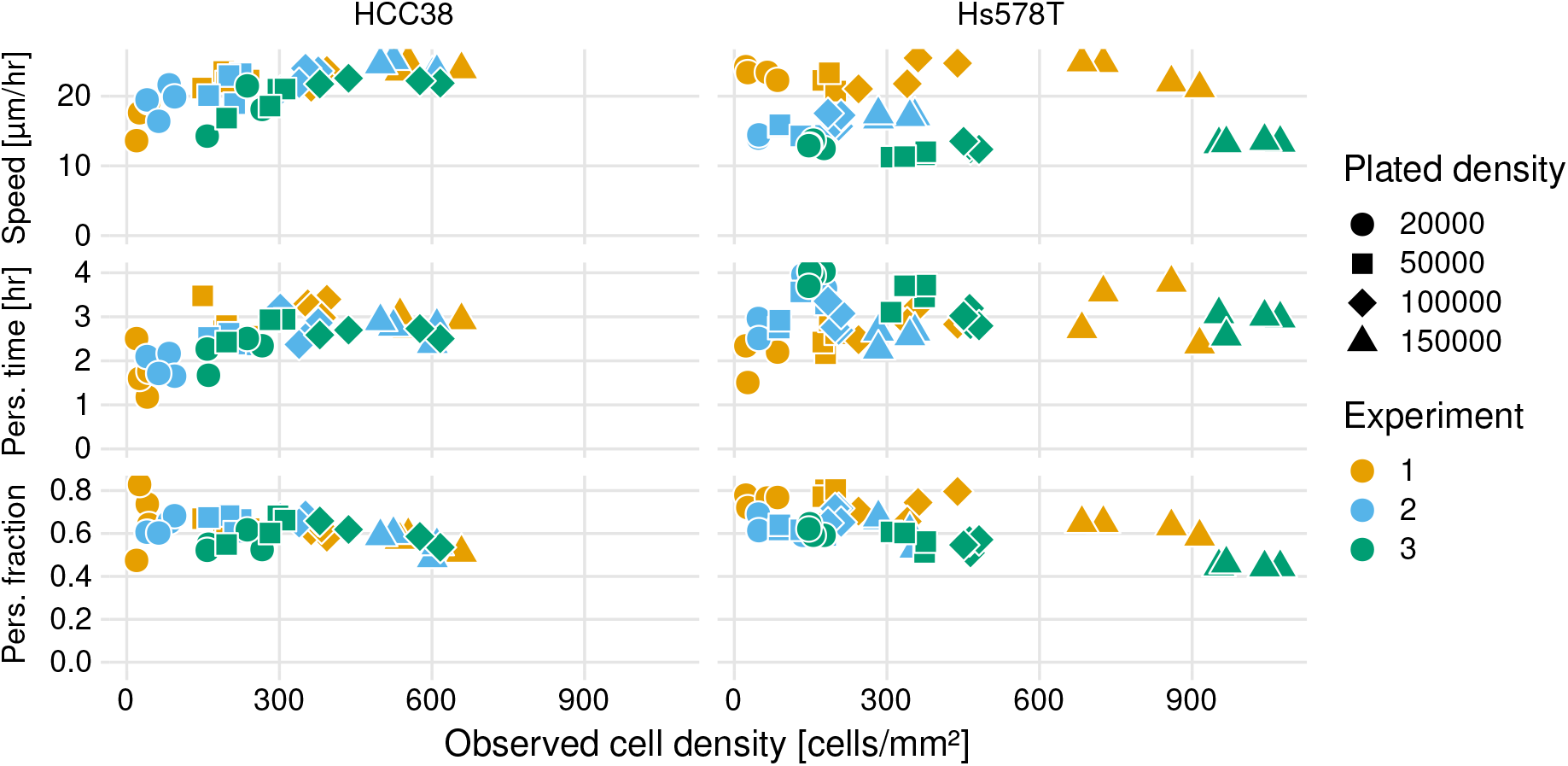
Speed, persistence time, and persistence fraction vs. observed cell density. Estimated parameter values were based on fitting exponential decay of the DAC (see Methods).

**Figure S7:**
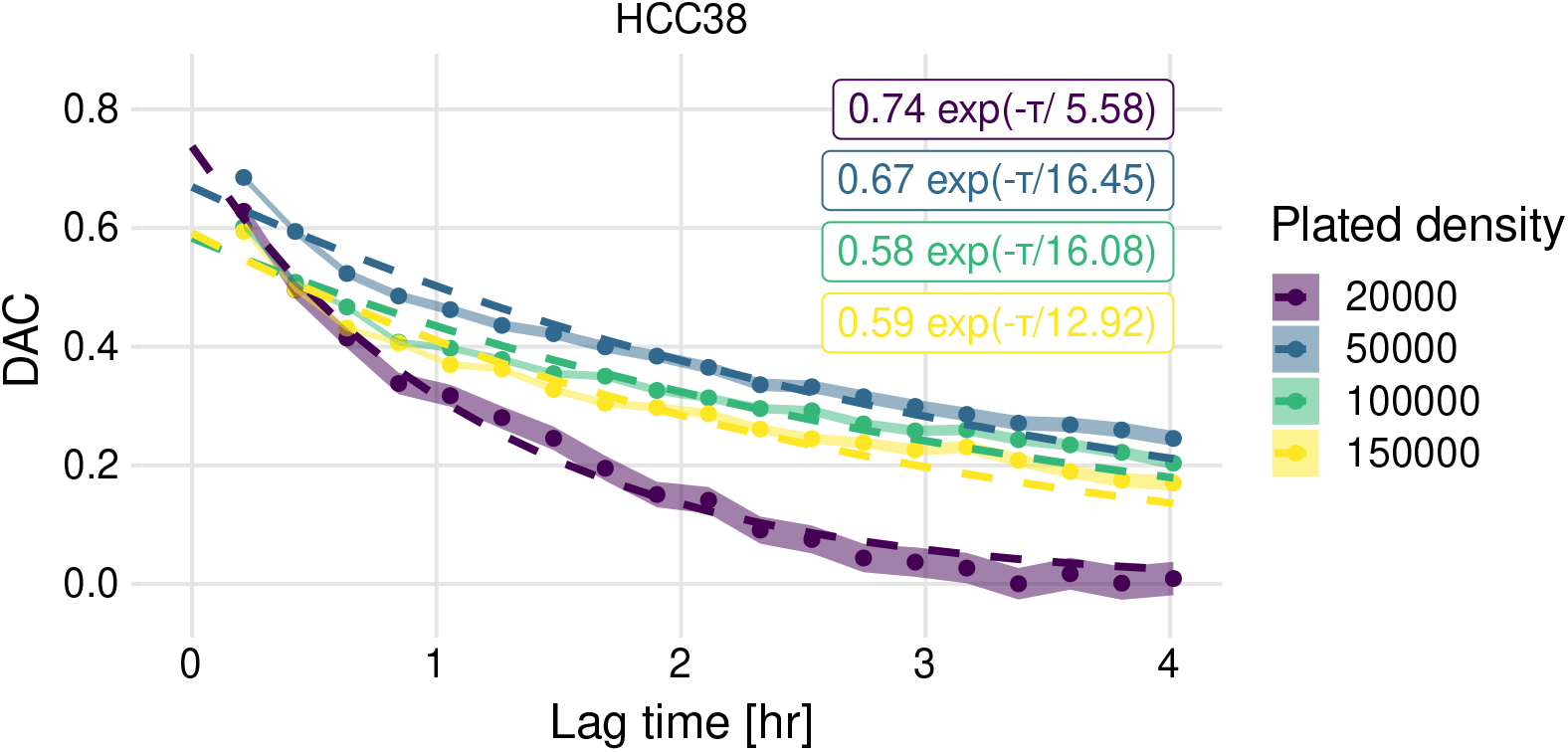
Example DAC fits for all densities of HCC38. The long-run persistence time is fitted well, but the persistent fraction (which is the intersect with the y-axis) is fitted quite poorly.

**Figure S8:**
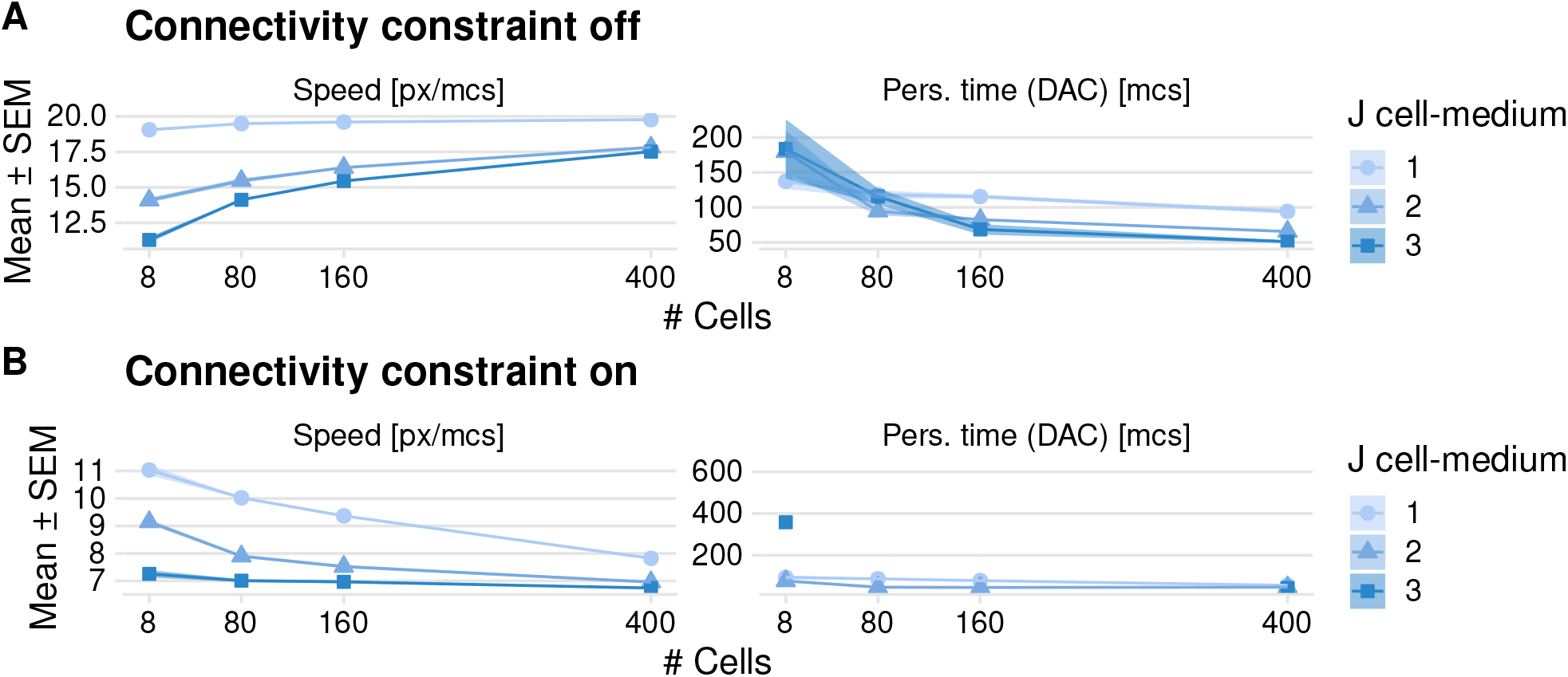
Influence of the connectivity constraint on cellular motility simulated with the basic persistence model using the Morpheus PersistentMotion plugin.

**Figure S9:**
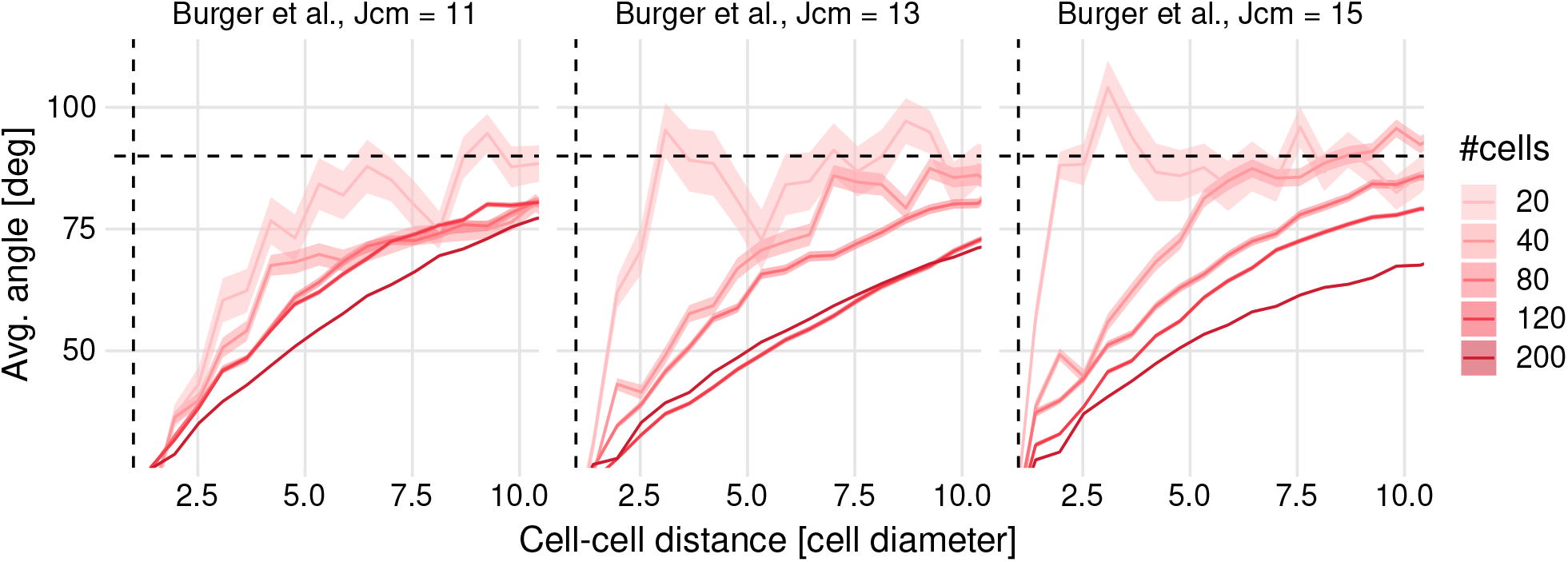
Influence of surface energy *J*_cell,med_ and of cell density on stream formation. Results are shown for the proposed persistence model with strong pseudopod coordination.

### Videos

**Video S1**. Experimental videos, same wells as in Fig. S1. Also available in higher quality at https://youtu.be/VFGVDyX_gI4.

**Video S2**. Persistence motion with basic persistence implemented in the PersistentMotion plugin in Morpheus. Rows are without (top) and with (bottom) connectivity constraint. *J*_cell,med_ = 3. Also available in higher quality at https://youtu.be/lzZJuFTNGC0.

**Video S3**. Actin protrusion plugin (Niculescu, Textor, and Boer, 2015). *J*_cell,med_ = 4. Also available in higher quality at https://youtu.be/TpjjyIsVVgU.

**Video S4**. Pseudopod-driven persistence model by Ariotti et al., 2012, implemented in Morpheus. *J*_cell,med_ = 5. Also available in higher quality at https://youtu.be/Wn4MP08AJHo.

**Video S5**. Proposed persistence model with strong pseudopod coordination. *J*_cell,med_ = 15. Also available in higher quality at https://youtu.be/7tpup52ERgo.

## Notes

https://doi.org/10.5281/zenodo.5484491

https://doi.org/10.5281/zenodo.5607734

### Competing Interest Statement

The authors have declared no competing interest.

## References

Aceto, Nicola et al. (Aug. 2014). “Circulating tumor cell clusters are oligoclonal precursors of breast cancer metastasis.” en. In: Cell 158.5, pp. 1110–1122. issn: 0092-8674, 0092-8674. doi: 10.1016/j.cell.2014.07.013.

Alamoudi, Emad et al. (n.d.). FitMultiCell: An Integrated Platform for Data-Driven Modeling of Multi-Cellular Processes. https://fitmulticell.gitlab.io/. Accessed: 2021-8-11.

Angelini, Thomas E et al. (Mar. 2011). “Glass-like dynamics of collective cell migration.” en. In: Proceedings of the National Academy of Sciences of the United States of America 108.12, pp. 4714–4719. issn: 0027-8424, 0027-8424. doi: 10.1073/pnas.1010059108.

Ariotti, Silvia et al. (Nov. 2012). “Tissue-resident memory CD8+ T cells continuously patrol skin epithelia to quickly recognize local antigen.” en. In: Proceedings of the National Academy of Sciences of the United States of America 109.48, pp. 19739–19744. issn: 0027-8424, 0027-8424. doi: 10.1073/pnas.1208927109.

Baddeley, Adrian, Ege Rubak, and Rolf Turner (Nov. 2015). Spatial Point Patterns: Methodology and Applications with R. en. CRC Press. isbn: 9781482210217.

Beerling, Evelyne et al. (Nov. 2016). “Intravital characterization of tumor cell migration in pancreatic cancer.” en. In: Intravital 5.3, e1261773. issn: 2165-9087, 2165-9087. doi: 10.1080/21659087.2016.1261773.

Beltman, Joost B, Athanasius F M Marée, and Rob J de Boer (Nov. 2009). “Analysing immune cell migration.” en. In: Nature reviews. Immunology 9.11, pp. 789–798. issn: 1474-1733, 1474-1733. doi: 10.1038/nri2638.

Beltman, Joost B, Athanasius FM Marée, Jennifer N Lynch, et al. (Apr. 2007). “Lymph node topology dictates T cell migration behavior.” en. In: The Journal of experimental medicine 204.4, pp. 771–780. issn: 0022-1007. doi: 10.1084/jem.20061278.

Bergert, Martin et al. (Sept. 2012). “Cell mechanics control rapid transitions between blebs and lamellipodia during migration.” en. In: Proceedings of the National Academy of Sciences of the United States of America 109.36, pp. 14434–14439. issn: 0027-8424, 0027-8424. doi: 10.1073/pnas.1207968109.

Bocci, Federico, Mohit Kumar Jolly, and José Nelson Onuchic (Nov. 2019). “A Biophysical Model Uncovers the Size Distribution of Migrating Cell Clusters across Cancer Types.” en. In: Cancer research 79.21, pp. 5527–5535. issn: 0008-5472, 0008-5472. doi: 10.1158/0008-5472.CAN-19-1726.

Brabletz, T et al. (2018). “EMT in cancer.” In: Nature reviews. Cancer. issn: 1474-175X.

Burger, Gerhard (Apr. 2021). burgerga/mdftracks. doi: 10.5281/zenodo.4692672.

Burger, Gerhard and Joost Beltman (Sept. 2021). morpheus-pseudopodia: A pseudopod-driven persistence model in Morpheus. doi: 10.5281/zenodo.5484491.

Burger, Gerhard, Muriel Heldring, and Steven Wink (Apr. 2021). burgerga/CPTrackR. doi: 10.5281/zenodo.4725473.

Buttenschön, Andreas and Leah Edelstein-Keshet (Dec. 2020). “Bridging from single to collective cell migration: A review of models and links to experiments.” en. In: PLoS computational biology 16.12, e1008411. issn: 1553-734X, 1553-7358. doi: 10.1371/journal.pcbi.1008411.

Carpenter, Anne E et al. (Oct. 2006). “CellProfiler: image analysis software for identifying and quantifying cell phenotypes.” en. In: Genome biology 7.10, R100. issn: 1465-6906. doi: 10.1186/gb-2006-7-10-r100.

Cordelières Fabrice P et al. (Nov. 2013). “Automated cell tracking and analysis in phase-contrast videos (iTrack4U): development of Java software based on combined mean-shift processes.” en. In: PloS one 8.11, e81266. issn: 1932-6203. doi: 10.1371/journal.pone.0081266.

Corey, David M, William P Dunlap, and Michael J Burke (July 1998). “Averaging Correlations: Expected Values and Bias in Combined Pearson rs and Fisher’s z Transformations.” In: The Journal of general psychology 125.3, pp. 245–261. issn: 0022-1309. doi: 10.1080/00221309809595548.

Fares, Jawad et al. (Mar. 2020). “Molecular principles of metastasis: a hallmark of cancer revisited.” en. In: Signal transduction and targeted therapy 5.1, p. 28. issn: 2059-3635. doi: 10.1038/s41392-020-0134-x.

Fougner, Christian et al. (Apr. 2020). “Re-definition of claudin-low as a breast cancer phenotype.” en. In: Nature communications 11.1, p. 1787. issn: 2041-1723. doi: 10.1038/s41467-020-15574-5.

Friedl, Peter et al. (Aug. 2012). “Classifying collective cancer cell invasion.” en. In: Nature cell biology 14.8, pp. 777–783. issn: 1465-7392, 1465-7392. doi: 10.1038/ncb2548.

Gallaher, Jill A, Joel S Brown, and Alexander R A Anderson (Feb. 2019). “The impact of proliferation-migration tradeoffs on phenotypic evolution in cancer.” en. In: Scientific reports 9.1, p. 2425. issn: 2045-2322. doi: 10.1038/s41598-019-39636-x.

Glazier, J A and F Graner (Mar. 1993). “Simulation of the differential adhesion driven rearrangement of biological cells.” en. In: Physical review. E, Statistical physics, plasmas, fluids, and related interdisciplinary topics 47.3, pp. 2128–2154. issn: 1063-651X. doi: 10.1103/PhysRevE.47.2128.

Gorelik, Roman and Alexis Gautreau (Aug. 2014). “Quantitative and unbiased analysis of directional persistence in cell migration.” en. In: Nature protocols 9.8, pp. 1931–1943. issn: 1754-2189, 1754-2189. doi: 10.1038/nprot.2014.131.

Graner, F and JA Glazier (Sept. 1992). “Simulation of biological cell sorting using a two-dimensional extended Potts model.” en. In: Physical review letters 69.13, pp. 2013–2016. issn: 0031-9007, 0031-9007. doi: 10.1103/PhysRevLett.69.2013.

Guillemot, J C et al. (Oct. 2001). “Ep-CAM transfection in thymic epithelial cell lines triggers the formation of dynamic actin-rich protrusions involved in the organization of epithelial cell layers.” en. In: Histochemistry and cell biology 116.4, pp. 371–378. issn: 0948-6143. doi: 10.1007/s004180100329.

Herschkowitz, Jason I et al. (2007). “Identification of conserved gene expression features between murine mammary carcinoma models and human breast tumors.” en. In: Genome biology 8.5, R76. issn: 1465-6906. doi: 10.1186/gb-2007-8-5-r76.

Hollestelle, Antoinette et al. (July 2010). “Four human breast cancer cell lines with biallelic inactivating alpha-catenin gene mutations.” en. In: Breast cancer research and treatment 122.1, pp. 125–133. issn: 0167-6806, 0167-6806. doi: 10.1007/s10549-009-0545-4.

Huang, Bin et al. (Dec. 2015). “Modeling the Transitions between Collective and Solitary Migration Phenotypes in Cancer Metastasis.” en. In: Scientific reports 5, p. 17379. issn: 2045-2322. doi: 10.1038/srep17379.

Huth, Johannes et al. (Apr. 2010). “Significantly improved precision of cell migration analysis in time-lapse video microscopy through use of a fully automated tracking system.” en. In: BMC cell biology 11, p. 24. issn: 1471-2121. doi: 10.1186/1471-2121-11-24.

Jacquemet, Guillaume et al. (Dec. 2016). “L-type calcium channels regulate filopodia stability and cancer cell invasion downstream of integrin signalling.” en. In: Nature communications 7, p. 13297. issn: 2041-1723. doi: 10.1038/ncomms13297.

Jayatilaka, Hasini et al. (May 2017). “Synergistic IL-6 and IL-8 paracrine signalling pathway infers a strategy to inhibit tumour cell migration.” en. In: Nature communications 8, p. 15584. issn: 2041-1723. doi: 10.1038/ncomms15584.

Kao, Jessica et al. (July 2009). “Molecular profiling of breast cancer cell lines defines relevant tumor models and provides a resource for cancer gene discovery.” en. In: PloS one 4.7, e6146. issn: 1932-6203. doi: 10.1371/journal.pone.0006146.

Kim, Donghwa et al. (Nov. 2019). “Antitumor Activity of Vanicoside B Isolated from Persicaria dissitiflora by Targeting CDK8 in Triple-Negative Breast Cancer Cells.” en. In: Journal of natural products 82.11, pp. 3140–3149. issn: 0163-3864, 0163-3864. doi: 10.1021/acs.jnatprod.9b00720.

Klijn, Christiaan et al. (Mar. 2015). “A comprehensive transcriptional portrait of human cancer cell lines.” en. In: Nature biotechnology 33.3, pp. 306–312. issn: 1087-0156, 1087-0156. doi: 10.1038/nbt.3080.

Kluyver, Thomas et al. (2016). “Jupyter Notebooks – a publishing format for reproducible computational workflows.” In: Positioning and Power in Academic Publishing: Players, Agents and Agendas. IOS Press, pp. 87–90. doi: 10.3233/978-1-61499-649-1-87.

Koedoot, Esmee et al. (Mar. 2021). “Differential reprogramming of breast cancer subtypes in 3D cultures and implications for sensitivity to targeted therapy.” en. In: Scientific reports 11.1, p. 7259. issn: 2045-2322. doi: 10.1038/s41598-021-86664-7.

Kramer, Nina et al. (Jan. 2013). “In vitro cell migration and invasion assays.” en. In: Mutation research 752.1, pp. 10–24. issn: 0027-5107. doi: 10.1016/j.mrrev.2012.08.001.

Le Dévédec, Sylvia E (Oct. 2021). HCC38 and MDA-MB-231 Triple-Negative Breast Cancer time lapses at different densities. doi: 10.5281/zenodo.5607734.

Litvinov, S V et al. (Dec. 1997). “Epithelial cell adhesion molecule (Ep-CAM) modulates cell-cell interactions mediated by classic cadherins.” en. In: The Journal of cell biology 139.5, pp. 1337–1348. issn: 0021-9525.

Liu, Yan-Jun et al. (Feb. 2015). “Confinement and low adhesion induce fast amoeboid migration of slow mesenchymal cells.” en. In: Cell 160.4, pp. 659–672. issn: 0092-8674, 0092-8674. doi: 10.1016/j.cell.2015.01.007.

Maiuri, Paolo et al. (Apr. 2015). “Actin flows mediate a universal coupling between cell speed and cell persistence.” en. In: Cell 161.2, pp. 374–386. issn: 0092-8674, 0092-8674. doi: 10.1016/j.cell.2015.01.056.

Mayor, Roberto and Sandrine Etienne-Manneville (Feb. 2016). “The front and rear of collective cell migration.” en. In: Nature reviews. Molecular cell biology 17.2, pp. 97–109. issn: 1471-0072, 1471-0072. doi: 10.1038/nrm.2015.14.

McCann, Colin P et al. (May 2010). “Cell speed, persistence and information transmission during signal relay and collective migration.” en. In: Journal of cell science 123.Pt 10, pp. 1724–1731. issn: 0021-9533, 0021-9533. doi: 10.1242/jcs.060137.

McCann, Kelly E, Sara A Hurvitz, and Nicholas McAndrew (July 2019). “Advances in Targeted Therapies for Triple-Negative Breast Cancer.” en. In: Drugs 79.11, pp. 1217–1230. issn: 0012-6667, 0012-6667. doi: 10.1007/s40265-019-01155-4.

Meijering, Erik, Oleh Dzyubachyk, and Ihor Smal (2012). “Methods for cell and particle tracking.” en. In: Methods in enzymology 504.February, pp. 183–200. issn: 0076-6879, 0076-6879. doi: 10.1016/B978-0-12-391857-4.00009-4.

Merks, Roeland M H et al. (Jan. 2006). “Cell elongation is key to in silico replication of in vitro vasculogenesis and subsequent remodeling.” en. In: Developmental biology 289.1, pp. 44–54. issn: 0012-1606. doi: 10.1016/j.ydbio.2005.10.003.

Morpheus (n.d.). Morpheus - TU Dresden. https://morpheus.gitlab.io/. Accessed: 2018-9-27.

Nair, Nishanth Ulhas et al. (July 2019). “Migration rather than proliferation transcriptomic signatures are strongly associated with breast cancer patient survival.” en. In: Scientific reports 9.1, p. 10989. issn: 2045-2322. doi: 10.1038/s41598-019-47440-w.

Neve, Richard M et al. (Dec. 2006). “A collection of breast cancer cell lines for the study of functionally distinct cancer subtypes.” en. In: Cancer cell 10.6, pp. 515–527. issn: 1535-6108. doi: 10.1016/j.ccr.2006.10.008.

Niculescu, Ioana, Johannes Textor, and Rob J de Boer (Oct. 2015). “Crawling and Gliding: A Computational Model for Shape-Driven Cell Migration.” en. In: PLoS computational biology 11.10, e1004280. issn: 1553-734X, 1553-7358. doi: 10.1371/journal.pcbi.1004280.

Parker, Joel S et al. (Mar. 2009). “Supervised risk predictor of breast cancer based on intrinsic subtypes.” en. In: Journal of clinical oncology: official journal of the American Society of Clinical Oncology 27.8, pp. 1160–1167. issn: 0732-183X, 1527-7755. doi: 10.1200/JCO.2008.18.1370.

Pau, Grégoire et al. (Apr. 2010). “EBImage–an R package for image processing with applications to cellular phenotypes.” en. In: Bioinformatics 26.7, pp. 979–981. issn: 1367-4803, 1367-4803. doi: 10.1093/bioinformatics/btq046.

Perou, C M et al. (Aug. 2000). “Molecular portraits of human breast tumours.” en. In: Nature 406.6797, pp. 747–752. issn: 0028-0836. doi: 10.1038/35021093.

Pijuan, Jordi et al. (June 2019). “In vitro Cell Migration, Invasion, and Adhesion Assays: From Cell Imaging to Data Analysis.” en. In: Frontiers in cell and developmental biology 7, p. 107. issn: 2296-634X. doi: 10.3389/fcell.2019.00107.

Prat, Aleix, Olga Karginova, et al. (Nov. 2013). “Characterization of cell lines derived from breast cancers and normal mammary tissues for the study of the intrinsic molecular subtypes.” en. In: Breast cancer research and treatment 142.2, pp. 237–255. issn: 0167-6806, 0167-6806. doi: 10.1007/s10549-013-2743-3.

Prat, Aleix, Joel S Parker, et al. (Sept. 2010). “Phenotypic and molecular characterization of the claudin-low intrinsic subtype of breast cancer.” en. In: Breast cancer research: BCR 12.5, R68. issn: 1465-5411, 1465-5411X. doi: 10.1186/bcr2635.

R Core Team (2018). R: A Language and Environment for Statistical Computing. Vienna, Austria.

Ribeiro, Ana Sofia and Joana Paredes (2014). “P-Cadherin Linking Breast Cancer Stem Cells and Invasion: A Promising Marker to Identify an “Intermediate/Metastable” EMT State.” en. In: Frontiers in oncology 4, p. 371. issn: 2234-943X. doi: 10.3389/fonc.2014.00371.

Ripley, B D (1977). “Modelling Spatial Patterns.” In: Journal of the Royal Statistical Society. Series B, Statistical methodology 39.2, pp. 172–212. issn: 1369-7412, 1369-7412.

Roosmalen Wies van et al. (2011). “Functional Screening with a Live Cell Imaging-Based Random Cell Migration Assay.” In: Cell Migration: Developmental Methods and Protocols. Ed. by Claire M Wells and Maddy Parsons. Totowa, NJ: Humana Press, pp. 435–448. isbn: 9781617792076. doi: 10.1007/978-1-61779-207-6/_29.

Rørth, Pernille (2009). “Collective cell migration.” en. In: Annual review of cell and developmental biology 25, pp. 407–429. issn: 1081-0706, 1081-0706. doi: 10.1146/annurev.cellbio.042308.113231.

RStudio Team (2016). RStudio: Integrated Development Environment for R. Boston, MA.

Rueden, Curtis T et al. (Nov. 2017). “ImageJ2: ImageJ for the next generation of scientific image data.” en. In: BMC bioinformatics 18.1, p. 529. issn: 1471-2105. doi: 10.1186/s12859-017-1934-z.

Schälte, Yannik et al. (July 2021). ICB-DCM/pyABC: pyABC 0.11.0. doi: 10.5281/zenodo.5149808.

Schindelin, Johannes et al. (June 2012). “Fiji: an open-source platform for biological-image analysis.” en. In: Nature methods 9.7, pp. 676–682. issn: 1548-7091, 1548-7091. doi: 10.1038/nmeth.2019.

Schindler, Daniel et al. (Aug. 2021). “Analysis of protrusion dynamics in amoeboid cell motility by means of regularized contour flows.” en. In: PLoS computational biology 17.8, e1009268. issn: 1553-734X, 1553-7358. doi: 10.1371/journal.pcbi.1009268.

Scianna, Marco and Luigi Preziosi (Mar. 2021). “A Cellular Potts Model for Analyzing Cell Migration across Constraining Pillar Arrays.” en. In: Axioms 10.1, p. 32. doi: 10.3390/axioms10010032.

Sørlie, T et al. (Sept. 2001). “Gene expression patterns of breast carcinomas distinguish tumor subclasses with clinical implications.” en. In: Proceedings of the National Academy of Sciences of the United States of America 98.19, pp. 10869–10874. issn: 0027-8424. doi: 10.1073/pnas.191367098.

Stanley, A J et al. (Dec. 1995). “Effect of cell density on the expression of adhesion molecules and modulation by cytokines.” en. In: Cytometry 21.4, pp. 338–343. issn: 0196-4763. doi: 10.1002/cyto.990210405.

Starruß, Jörn et al. (May 2014). “Morpheus: a user-friendly modeling environment for multiscale and multicellular systems biology.” en. In: Bioinformatics 30.9, pp. 1331–1332. issn: 1367-4803, 1367-4803. doi: 10.1093/bioinformatics/btt772.

Stramer, Brian and Roberto Mayor (Jan. 2017). “Mechanisms and in vivo functions of contact inhibition of locomotion.” en. In: Nature reviews. Molecular cell biology 18.1, pp. 43–55. issn: 1471-0072, 1471-0072. doi: 10.1038/nrm.2016.118.

Stuelten, Christina H, Carole A Parent, and Denise J Montell (May 2018). “Cell motility in cancer invasion and metastasis: insights from simple model organisms.” en. In: Nature reviews. Cancer 18.5, pp. 296–312. issn: 1474-175X, 1474-1768. doi: 10.1038/nrc.2018.15.

Suhail, Yasir et al. (Aug. 2019). “Systems Biology of Cancer Metastasis.” en. In: Cell systems 9.2, pp. 109–127. issn: 2405-4720, 2405-4720. doi: 10.1016/j.cels.2019.07.003.

Svensson, Carl-Magnus et al. (Mar. 2018). “Untangling cell tracks: Quantifying cell migration by time lapse image data analysis.” en. In: Cytometry. Part A: the journal of the International Society for Analytical Cytology 93.3, pp. 357–370. issn: 1552-4922, 1552-4922. doi: 10.1002/cyto.a.23249.

Szabó, A et al. (Nov. 2010). “Collective cell motion in endothelial monolayers.” en. In: Physical biology 7.4, p. 046007. issn: 1478-3967, 1478-3967. doi: 10.1088/1478-3975/7/4/046007.

Szabó, András, Manuela Melchionda, et al. (June 2016). “In vivo confinement promotes collective migration of neural crest cells.” en. In: The Journal of cell biology 213.5, pp. 543–555. issn: 0021-9525, 0021-9525. doi: 10.1083/jcb.201602083.

Szabó, András and Roeland M H Merks (Apr. 2013). “Cellular potts modeling of tumor growth, tumor invasion, and tumor evolution.” en. In: Frontiers in oncology 3, p. 87. issn: 2234-943X. doi: 10.3389/fonc.2013.00087.

Te Boekhorst Veronika, Luigi Preziosi, and Peter Friedl (Oct. 2016). “Plasticity of Cell Migration In Vivo and In Silico.” en. In: Annual review of cell and developmental biology 32, pp. 491–526. issn: 1081-0706, 1081-0706. doi: 10.1146/annurev-cellbio-111315-125201.

Van Haastert, Peter J M (Feb. 2011). “How cells use pseudopods for persistent movement and navigation.” en. In: Science signaling 4.159, e6. issn: 1937-9145, 1937-9145. doi: 10.1126/scisignal.2001708.

Van Liedekerke, P et al. (Dec. 2015). “Simulating tissue mechanics with agent-based models: concepts, perspectives and some novel results.” In: Computational Particle Mechanics 2.4, pp. 401–444. issn: 2196-4386. doi: 10.1007/s40571-015-0082-3.

Vedel, Søren et al. (Jan. 2013). “Migration of cells in a social context.” en. In: Proceedings of the National Academy of Sciences of the United States of America 110.1, pp. 129–134. issn: 0027-8424, 0027-8424. doi: 10.1073/pnas.1204291110.

Viale, G (Sept. 2012). “The current state of breast cancer classification.” en. In: Annals of oncology: official journal of the European Society for Medical Oncology /ESMO 23 Suppl 10, pp. x207–10. issn: 0923-7534, 0923-7534. doi: 10.1093/annonc/mds326.

Waclaw, Bartlomiej et al. (Sept. 2015). “A spatial model predicts that dispersal and cell turnover limit intratumour heterogeneity.” en. In: Nature 525.7568, pp. 261–264. issn: 0028-0836, 0028-0836. doi: 10.1038/nature14971.

WCRF (Jan. 2021). Breast cancer statistics. en. https://www.wcrf.org/dietandcancer/breast-cancer-statistics/. Accessed: 2021-8-10.

Wickham, Hadley (2017). tidyverse: Easily Install and Load the ‘Tidyverse’.

Wink, Steven (2015). “Systems microscopy to unravel cellular stress response signalling in drug induced liver injury.” PhD thesis. Leiden University.

Wink, Steven and Gerhard Burger (Sept. 2021). H5CellProfiler. doi: 10.5281/zenodo.5484432.

Winter, Manon J et al. (Apr. 2003). “Expression of Ep-CAM shifts the state of cadherin-mediated adhesions from strong to weak.” en. In: Experimental cell research 285.1, pp. 50–58. issn: 0014-4827. doi: 10.1016/S0014-4827(02)00045-9.

Wortel, Inge M N, Katharina Dannenberg, et al. (June 2019). “CelltrackR: an R package for fast and flexible analysis of immune cell migration data.” en.

Wortel, Inge M N, Ioana Niculescu, et al. (July 2021). “Local actin dynamics couple speed and persistence in a cellular Potts model of cell migration.” en. In: Biophysical journal 120.13, pp. 2609–2622. issn: 0006-3495, 0006-3495. doi: 10.1016/j.bpj.2021.04.036.

Wu, Pei-Hsun et al. (Mar. 2014). “Three-dimensional cell migration does not follow a random walk.” en. In: Proceedings of the National Academy of Sciences of the United States of America 111.11, pp. 3949–3954. issn: 0027-8424, 0027-8424. doi: 10.1073/pnas.1318967111.

Yamamoto, Mizuki et al. (June 2017). “Intratumoral bidirectional transitions between epithelial and mesenchymal cells in triple-negative breast cancer.” en. In: Cancer science 108.6, pp. 1210–1222. issn: 1347-9032, 1347-9032. doi: 10.1111/cas.13246.

Yan, Kuan and Fons J Verbeek (Oct. 2012). “Segmentation for High-Throughput Image Analysis: Watershed Masked Clustering.”en. In: Leveraging Applications of Formal Methods, Verification and Validation. Applications and Case Studies. Ed. by Tiziana Margaria and Bernhard Steffen. Lecture Notes in Computer Science. Springer Berlin Heidelberg, pp. 25–41. isbn: 9783642340314. doi: 10.1007/978-3-642-34032-1_4.

